# Systematic mapping shows monitoring, evaluation, and engagement are needed to build an evidence base for rewilding

**DOI:** 10.64898/2026.01.02.697359

**Authors:** Nell J. Pates, James M. Bullock, Johan T. du Toit, Sahran Higgins, Nathalie Pettorelli, William D. Pearse

## Abstract

Rewilding is a landscape recovery approach that seeks to enhance functioning of degraded ecosystems by reinstating natural processes and is increasingly being adopted by organisations and landowners. Despite its popularity, there is no accepted way to assess rewilding’s impacts. This is due in part to inconsistent monitoring and evaluation. Here we report the results from a systematic map of the academic and grey literature on monitoring and evaluation in rewilding case studies. We find that researchers and practitioners are focused on different aspects of rewilding, and that limited recording of case study details and lack of robust experimental design make it difficult to pool and compare data. We identify challenges facing researchers and rewilding practitioners seeking to build a robust evidence base from which to evaluate rewilding outcomes. These challenges include resources as well as spatial and time constraints, which all limit the consistency of data gathering among projects. More comparisons across rewilding projects, better standardisation of data collection and data recording methods, and increased collaboration between researchers and practitioners could go some way to remedy this. We provide recommendations for planning and implementing studies of rewilded landscapes to aid monitoring and evaluation. These include the broad adoption of low-cost and scalable survey designs with greater focus on comparisons across sites, and continued effort towards open-source data storage.

## Introduction

Most of the planet’s land surface has been measurably impacted by human activity (Kennedy et al., 2019, Venter et al., 2016) with around a third estimated to be under intense pressure (Jones et al., 2018). These pressures degrade ecosystems (Williams et al., 2020), leading to increased extinction risk for species (Di Marco et al., 2018) and loss of ecosystem functions, and services for humans who rely on them (IPBES, 2019, Isbell et al., 2023). Given the intrinsic value of nature and its importance for securing human wellbeing, there is an urgent need not only to protect remaining areas of conservation value but also to recover damaged ecosystems. This need is reflected in recent roadmaps produced both in policy (UN Decade on Ecosystem Restoration, 2021-2030) and academic research (Dinerstein et al., 2020, Maron et al., 2020, Yang et al., 2020). Within this context, rewilding has emerged in recent years as a popular approach to reversing biodiversity loss and ecosystem degradation (Pettorelli and Bullock, 2023).

Much discussion has been given to establishing a universal definition for rewilding (Corlett, 2016, Pettorelli et al., 2018) and a degree of consensus seems to have emerged in recent years (Carver et al., 2025a). Rewilding seeks to enhance the functional properties of degraded ecosystems so they can become more self-sustaining, requiring minimal long-term management (Perino et al, 2019, Pettorelli et al, 2018). Developed since the 1990s, rewilding has been credited with injecting a degree of excitement and inspiration into landscape recovery (Wynne-Jones et al., 2020) and is now a mainstream strategy in several countries (Carver et al., 2025a) to the extent that organisations are producing practical guides and handbooks to support the initiation and maintenance of rewilding projects (IUCN Commission on Ecosystem Management: Carver et al, 2025b; Rewilding Europe: Rewilding Europe, 2025; WWF-UK: Dempsey, 2023).

Despite the growing interest, gaps remain in understanding the long-term outcomes of rewilding projects. These gaps are driven in part by rewilding’s appeal: there has been rapid adoption of rewilding, from vast landscape-level interventions to relatively tiny community projects, with an associated surge in research interest (Lorimer et al., 2015, Hart et al., 2023). However, monitoring and evaluation have not kept pace with implementation. A number of opinion and review pieces about rewilding have emerged (e.g., Egoh et al, 2021), but very few long-term, comparable studies, and this is a significant limitation to the study of rewilding (Hart et al., 2023). To determine whether rewilding can deliver on what it offers – transition of degraded landscapes to more self-sustaining ecosystems – in a way that is valuable to practitioners who may look for evidenced-based guidance to choose one course of action over another, we need to assess its effectiveness over time through monitoring and evaluation. Rewilding projects pose specific challenges for monitoring and evaluation for two main reasons. Firstly, as rewilding is a relatively young strategy, many projects could not have accrued long-term data sets even if effective monitoring were in place. Secondly, evaluation often requires an expected goal to compare progress against, but what is meant by ‘success’ is not necessarily clear-cut in rewilding. Projects may frame their terms for success more broadly than biodiversity enhancement, for example by referring to specific rewilding principles or their own individual objectives linked to their environmental and social context; in that context, goals may be short, medium, or long term.

This study aimed to assess the current state of literature on monitoring and evaluation in rewilding, while acknowledging the context of the challenges outlined above, to assess emerging patterns. We present a systematic map of monitoring and evaluation in rewilding case studies using both peer-reviewed and grey literature (literature from outside of traditional academic publishing, such as organisations’ reports and policy documents) to investigate how success is defined in rewilding, and how progress in rewilding projects is monitored and evaluated. We sought to establish what is being measured currently, where, and by whom. We compare approaches and metrics used by researchers and practitioners to look for areas of overlap and difference. Improved knowledge and understanding of what is working in current re-wilding projects is required to form a strong evidence base, which may assist re-wilding practitioners to identify and implement good practice. This will be critical in securing support and funding if rewilding strategies are to be implemented at scale to achieve their global potential.

## Methods

We conducted systematic mapping of academic and grey literature with a focus on monitoring and evaluation in rewilding case studies. Academic literature was defined as academic publications, including theses. Grey literature is a term that captures a broad range of materials, including that which is not typically peer reviewed, and not produced for commercial publication or wide distribution (Tillett and Newbold, 2006). The grey literature considered here was information and literature provided by, or available by public access from, organisations belonging to the Global Rewilding Alliance (GRA). A systematic map (also known as a scoping review, or evidence map) collates available information to identify gaps in the current understanding of a field (Arksey and O’Malley, 2005, Haddaway et al., 2020). Systematic maps differ from full systematic literature reviews (Munn et al., 2018), which are most useful when a body of detailed, quantitative, literature exists. Systematic mapping, on the other hand, is appropriate when the available information is sparse, complex and / or heterogeneous (James et al., 2016, Peters et al., 2021), as is the case in rewilding. Systematic maps aim to identify gaps in data, while summarising existing trends in a clear and accessible format (Arksey and O’Malley, 2005, James et al., 2016). Here we identified knowledge gaps and areas of overlap within and between the academic and grey data sets. The systematic approach employed here means that future attempts to understand what we have learned can build directly on this transparent and repeatable methodology.

### Key definitions

This systematic map is focused on monitoring and evaluation in re-wilding, and definitions were required to draw conclusions from the collected literature. **Monitoring** is the practice of gathering data over time. To be classified as a monitoring study here, academic papers had to include repeated measures over time – usually over several years. **Evaluation** is the process of assessing whether an action – in this case, rewilding – has achieved a desired outcome. To be classified as an evaluation study, academic papers had to make clear the aims of their intervention and whether the results of the study were in keeping with those aims. Re-wilding organisations were asked directly about their monitoring and evaluation practices, and organisations’ policy documents and websites were searched for information conforming with these definitions.

### Research questions

We investigated three research questions: (1) How is success in rewilding defined?; (2) How is progress towards success in rewilding projects monitored?; (3) How is progress towards success in rewilding projects evaluated?

These questions capture some known difficulties in studying rewilding. Firstly, the idea of success in any rewilding intervention is not necessarily clear cut, and desired rewilding outcomes may not even be specified (Law et al., 2017). Where there is a known intended outcome, it might be tied to target species, habitat structure, some level of ecological function or service, or may be based on, for example, a known pristine remnant, an estimate of a past state, or a hypothetical prediction (i.e. a projected novel system; Sinclair et al., 2018). We have provided a definition of rewilding to frame our study but, given that a universally agreed definition does not exist, we did not expect to find a consistent definition across case studies, and we did not seek to compare the definitions of success we found against any benchmark. Rather, we were interested in the content of each description of success, and how the two groups -researchers and practitioners -use terms to conceptualise what they aim to achieve through their studies and projects. Descriptions of success for case studies were not readily available within the academic literature as results are not framed in terms of the landscape of interest, but with reference to study objectives and hypotheses. To draw comparable data from the academic literature we extracted the study focus (the taxa or system of interest) of each paper as a proxy for a definition of success. This gave an indication of what aspects of landscape recovery are receiving the most attention. We may infer from this which groups are considered most important by the research community, but we acknowledge that this measure of ‘success’ is somewhat arbitrary and captures only one aspect of each study.

We also anticipated complexity in investigating the monitoring and evaluation research questions. Rewilding outcomes are challenging to measure. Firstly, because efforts to measure ecosystem-level impacts are likely to be very time- and resource-intensive. Secondly, because ecosystem recovery can be a long, slow process that is difficult to measure meaningfully in a short time frame (Corlett, 2019), practitioners may have decades- or centuries-long goals for their landscapes while researchers will typically be constrained by the time and resources available for them to plan and conduct a study. Thirdly, each rewilding case study is positioned within a specific socio-environmental context that will impact how researchers and practitioners approach it. Here, the aim was to build up a picture of how monitoring and evaluation strategies compare across the two strands of literature.

### Academic literature review

We followed a 5-stage review process based on systematic mapping tools described in James et al. (2016) and Levac et al. (2010) to analyse the academic literature: (1) preliminary activities, (2) search, (3) screen, (4) extract, (5) report (Figure 1). This clear, staged methodology brings consistency and rigor to the process mapping, allowing for evaluation and refinement at each stage.

**Figure 1.**
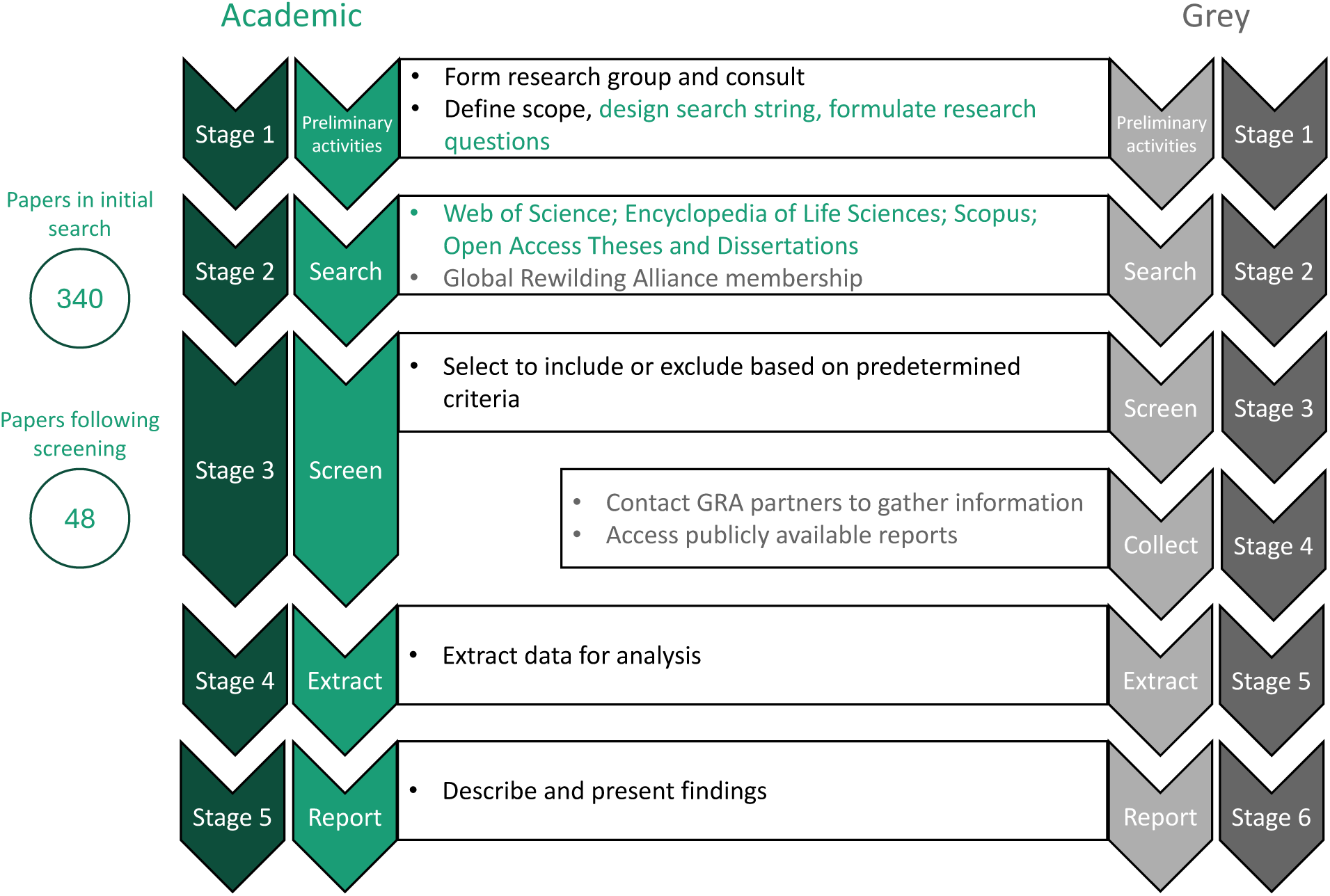
The 5-stage methodology used for review of academic literature, and 6-stage methodology used for grey literature to produce our systematic map. This approach is designed to be as transparent, unbiased, and – critically – repeatable as possible. It should be possible for later studies to follow the same steps and assess what progress has been made against our results. Although presented as a linear process for simplicity, the process is iterative (Levac et al., 2010). Black text describes steps for both academic and grey literature. Green text describes actions performed for academic literature only. Grey text describes actions performed for grey literature only.

#### (1) Preliminary activities

We determined the scope of this study, research questions, key definitions, and the search string. We also selected criteria for inclusion during the screening stage. Consultation with a research team is a valuable component in the framing of systematic reviews of this kind – particularly in reviews dealing with grey literature (Arksey and O’Malley, 2005) – providing a breadth of ideas that might otherwise be overlooked. The authors represent a range of expertise in rewilding, both as academics and practitioners. The academic consultation team (NP, JdT, JB) reviewed and provided feedback on early iterations of the scope, research questions, search string and criteria, as well as interpretation of results later in the process. SH provided a practitioner perspective from the results stage onwards.

##### Scope

The scope of this academic literature review was English-language academic publications, including theses, that describe monitoring and / or evaluation of re-wilding case studies. Although a single language focus is a possible source of bias in this study (Amano et al, 2016, Nuñez and Amano, 2021), this decision reflects the fact that ‘rewilding’ is still a fluid term and many case studies that may be classified as rewilding under some definitions may be excluded under others. The aim of this study was not to impose our own definition on studies or organisations, but to investigate how different groups conceive and measure rewilding in their case studies. We therefore searched for studies that explicitly described rewilding case studies, without applying our own definition as a requirement for inclusion. We feel it is important to comprehensively survey the state of rewilding, rather than excluding groups based on any particular definition of rewilding; this ensures learnings derived from this study build on all the work done in this space.

##### Search string

The search string used for the literature search was: *“rewild* AND (evaluat* OR monitor*)”*. Our search string was validated through scoping – testing and adjusting the search string to ensure outputs were appropriately sensitive and specific (James et al., 2016). This search string will not have captured projects that may share the same aims and methods as those that associate themselves explicitly with ‘rewilding’ but use different terminology. Similarly, studies whose methods may meet the definitions of monitoring and evaluation outlined above but that used ‘measure’ or ‘assess’, for example, may not have been captured. It is, therefore, somewhat exclusive of candidate case studies. However, this degree of rigor limits the requirement for significant interpretation at the screening stage and retains tight focus on the phenomenon known as ‘rewilding’.

#### (2) Search

We conducted the academic literature search on 6 November 2024 (Figure 1), using our search string across multiple databases (Web of Science, Scopus, PubMed, and Open Access Theses and Dissertations) in an effort to cast the net widely for available literature (Haddaway et al., 2015b).

#### (3) Screen

We downloaded titles and abstracts of all search results. If abstracts were missing, we added them to the dataset manually before screening. We screened the results of our literature search in several stages (see supplementary Table S1.1). First, we pooled all results and removed duplicates. Then we used the R package metagear (Lajeunesse, 2016) to screen entries by title and abstract for inclusion using pre-determined criteria (code available at DOI: 10.5281/zenodo.18131011). The final stage of screening was full paper review, where we read and compared each paper against a further set of criteria (see supplementary information section 1.1 for criteria).

#### (4) Extract

We extracted three main categories of data from each paper: bibliographic information (authors, year of publication, title); case study information (location, study area etc); and research question information (focus study system, methods of monitoring etc). Some interpretation of data was necessary to allow coding (assigning extracted metadata to predetermined categories to allow comparison) of the extracted information (James et al., 2016). Full detail is provided in supplementary information 1 (section 1.2, Table S1.2).

#### (5) Report

A narrative synthesis is the principal output of systematic mapping (James et al., 2016). This paper is the culmination of our review, presenting our results and reporting summary statistics to describe the current state of literature on this topic.

### Grey literature review

We slightly adapted the process described for the academic literature review to collect and analyse grey literature on rewilding case studies. The grey literature review was completed in 6 stages: preliminary activities, search, screen, collect, extract, report (Figure 1).

#### (1) Preliminary activities

Other reviews that have included grey literature have relied on existing professional networks to gather documentation (e.g. Van Klink and WallisDeVries (2018), Langridge et al. (2021)). We opted to follow a transparent and repeatable methodology that does not rely on existing relationships. We identified the Global Rewilding Alliance (2024) – a global partnership that allows free, voluntary membership of organisations and individuals involved in rewilding – as a source for our grey literature search. The GRA maintains a list of alliance partner organisations, and we used this to search and survey. This method will remain available to future reviews, even as the GRA membership changes over time, for as long as the GRA exists.

##### Scope

The scope of the grey literature review was information and literature provided by, and / or publicly available from, organisations belonging to the GRA. The request for direct communication from organisations was an important aspect of this study. Although not a traditional source of grey literature (Haddaway et al, 2015a), using email or interview responses to standard questions allowed us to maximise the available information. This was valuable because organisational documents and websites were not typically aligned to our exact research questions and were not comparable in the manner of academic research articles. In addition, the grey literature overall was very complex. Direct email responses and subsequent discussions where meetings were held frequently comprised very narrative responses and these had to be recorded carefully.

In our review of grey literature, we were not restricted to English language documents but searched for information across all relevant organisations regardless of language. In most cases, information was available in English either because it was produced in English, or because websites provided multiple language documents, or had inbuilt translation functions. In these cases, the English language version was used for analysis. Where it was not possible to access English language documents (three Spanish, three Portuguese, one Chinese, and one Dutch) we sought native speakers of these languages and explained the protocol to them to extract data for these organisations. We did not have access to a native Portuguese language speaker, so for these three organisations we used Google Translate.

#### (2) Search

The GRA membership list, as of November 2024, was the start point for our grey literature review (Global Rewilding Alliance, 2024).

#### (3) Screen

We checked the organisational websites of each GRA partner and selected 93 from a total of 194 by excluding partner organisations that were not directly involved with rewilding case studies, e.g. bookstores, magazines, or rewilding photographers.

#### (4) Collect

We collected data from the screened subset of GRA partners in two stages.

##### (i) Direct contact

We contacted organisations directly by email (see supplementary information 1, section 2.1.1) to elicit comparable responses about re-wilding organisations’ approaches to defining success in rewilding, monitoring and evaluating progress. The first round of emails was sent on 16 October 2024. A repeat was sent on 7 November 2024 and 23 organisations responded to the email: 16 provided detailed replies; 7 declined.

Representatives from four organisations requested to meet virtually to discuss their work: we met online and took notes.

##### (ii) Documents

We collected documents, either provided by respondents to the email approach, or publicly available through their websites. We collected documents for all organisations, regardless of whether they responded to the email approach.

Following data collection, eight organisations were removed from the data set. These were organisations nested within other GRA members that had already provided data, or organisations working to bring together rewilding resources or secure funding, rather than operating on the ground. The final data set therefore contains 85 re-wilding organisations.

#### (5) Extract

We extracted four main categories of data about each organisation: partner organisation detail (name of organisation, country and continent of operation); direct communication detail (whether or not a response was received and what detail or documents were included); case study information (area, start year of project); research question information (details related to success, monitoring and evaluation) see supplementary information 1 (section 2.2, Table S1.5). Within the research question information section, we assessed definitions of success as ‘specific’, ‘measurable’, ‘time-bound’, or any combination of the three, in line with the concept of SMART (specific, measurable, ambitious, realistic, and time-bound) goal setting (Maxwell et al., 2015) and recorded these to compare across the data set.

#### (6) Report

As above, this paper is the culmination of our review.

## Results

We performed a full review of 48 academic papers following screening. The papers were published between 2015 and 2024, and included case studies from 40 countries (see supplementary Figure S2.1A). We categorised papers as ‘monitoring’ (25%), ‘evaluation’ (40%), or ‘monitoring and evaluation’ (35%) using our definitions. The 85 rewilding organisations included in the grey literature review had rewilding projects in 106 countries (Figure S2.1B). We found documents including information for at least one of our research questions (success, monitoring, and / or evaluation) for 81 out of 85 organisations. This was supplemented with information from direct contact for the 16 organisations that replied to the email. Most rewilding organisations provided a definition of success (86%), but fewer outlined their monitoring (64%) or evaluation (52%) work. Study areas in academic case studies (academic literature) were typically smaller than the areas managed by rewilding practitioners (grey literature): the median area of academic case studies was 13.7km^2^ (interquartile range (IQR) = 57.2km^2^), compared with the median area in the grey literature of 309km^2^ (IQR = 9,756km^2^). Most academic case studies reported on one or two years of data (26 out of 48 studies, median duration = 1 year, IQR = 1). In the academic literature, 60% of case studies (n = 29) did not describe controls as part of their design. In the grey literature, two out of 85 rewilding organisations described controls implemented as part of their monitoring strategies.

### Question 1: How is success in rewilding defined?

We compared the study focus of academic case studies with definitions of success provided by rewilding organisations or found within their documents. There was some overlap between categories but there was a disconnect between the amount of attention given to categories common to both strands of literature. In the academic literature, most studies focused on mammals (35%) and plants (27%) (Figure 2A). In the grey literature, fewer definitions of success centred around species considerations, but these also showed a bias in favour of mammals (65% of species-centric definitions). Percentages related to these data refer to the proportion of categories, rather than studies or organisations. The most common definitions of success for rewilding organisations were tied to social considerations (23%), such as public engagement, empowering communities, providing jobs, developing tourism etc. (Figure 2B). Social considerations featured in 6% of academic papers. We assessed rewilding organisations’ definitions of success as ‘specific’, ‘measurable’, ‘time-bound’. Just 19% of organisations had definitions of success that we classified as specific, measurable, *and* time-bound, and 37 definitions did not meet any of the three criteria (including instances where no definition was provided).

**Figure 2.**
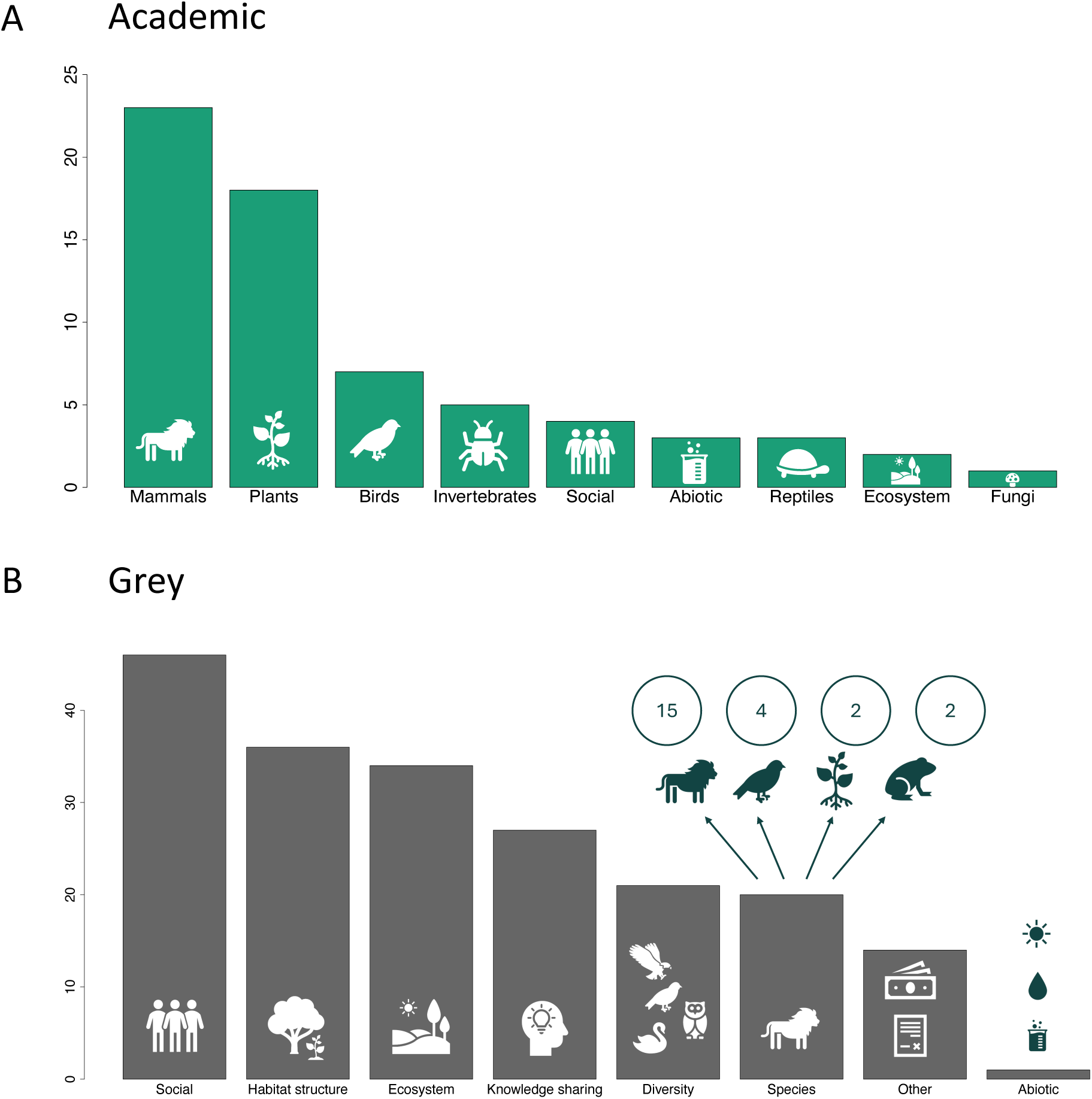
Comparing academic case studies’ focus study systems and rewilding organisations’ definitions of success reveals different priorities. **A.** Academic studies were assigned categories to understand which taxa received the most attention. Mammals and plants were the most commonly studied categories. **B.** Most practitioner definitions of success in rewilding projects related to social outcomes. The ‘other’ category for grey literature captures legal (e.g. secure formal protections), financial (e.g. become profitable), and definitions of success linked to policy targets (e.g. meet national and international biodiversity goals). Sums of bar plots are not the counts of academic papers reviewed or rewilding organisations contacted. Some academic studies were assigned two (n = 14) or three (n = 2) focus categories where they studied multiple taxonomic groups, for example. Of the 72 rewilding organisations for which definitions of success were included, 60 definitions spanned multiple categories, partly due to the long and narrative data available (2 categories: n = 24; 3 categories: n = 19; 4 categories: n = 9; 5 categories: n = 5; 6 categories n = 2, 8 categories n = 1). Full detail on each category and how these were assigned is provided in supplementary information 1.

### Question 2: How is progress towards success in rewilding projects monitored?

We extracted detail on the methods used in academic studies for both monitoring and evaluation and compared these with strategies used for monitoring by rewilding organisations. There was overlap between the two strands of literature in this research question. Fieldwork (direct observations through surveys, transects etc.) emerged as the most common method used to collect data for monitoring and evaluation (41% in academic literature, 39% in grey). Both strands also included wildlife tracking, remote sensing, experimental, consultation, and DNA methods in varying proportions (Figure 3). One category – ‘review’ – was present in the academic but not the grey literature, and an ‘aspirational’ category was included in the grey review to capture monitoring efforts that organisations hoped or planned to implement in the future.

**Figure 3.**
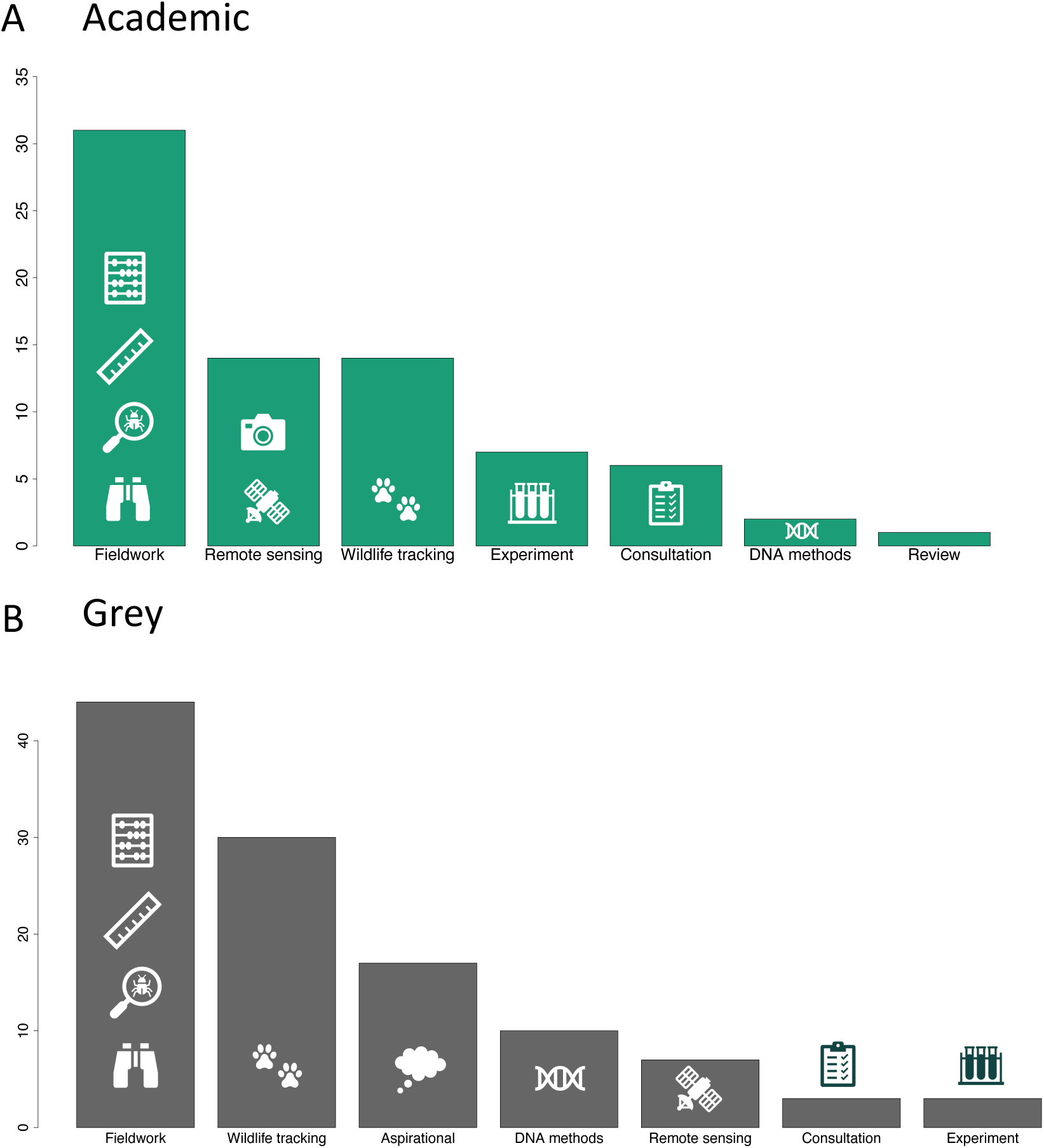
Fieldwork is the most common method described in both academic and grey literature to investigate rewilding progress. **A.** The methods described in each academic paper were coded and categorised. Multiple categories were assigned to studies that used more than one approach**. B.** Data was extracted from direct communication or analysis of grey literature, then coded following the same process as used for the academic literature. Full detail on each category and how these were assigned can be found in supplementary information 1.

### Question 3: How is progress towards success in rewilding projects evaluated?

We extracted detail on the metrics used in academic studies and by rewilding organisations to measure rewilding outcomes. For the academic literature, we extracted response variables from vertical plot axes, figure and table legends, and / or sub-headings of results sections and found 212 unique response variables. These were coded into categories for analysis. Grey literature was coded from whole sentences and paragraphs from direct contact or documentation. Comparison of categories revealed differences between the kinds of metrics used by researchers and practitioners to evaluate success in rewilding. Only three categories were common to both strands (diversity, species, and social metrics). Academic studies focused most on distribution metrics (abundance, density, etc -20%) and habitat structure (habitat classification, grass cover etc -22%) and also included metrics related to species behaviour, abiotic factors, and ecosystem metrics (Figure 4A). Four metrics (within two studies) were categorised as ‘other’: two from a study of vertebrate visitors to communal rhinoceros latrines which reported on latrine quality and size (Awasthi et al., 2024); and two from a study reporting the number of reintroduction projects and their objectives (Banasiak et al., 2021). In the grey literature, the most common evaluation metrics were ‘internal’, meaning metrics that are predominantly relevant to the GRA organisation itself (23%). Examples of internal category metrics included personnel hired or trained, size of organisation, number of projects planned and implemented. Other categories unique to the grey literature were external metrics (standards set by external bodies or organisations), measures of area, academic and financial considerations, and ‘aspirational’ to capture future intentions (Figure 4B).

**Figure 4.**
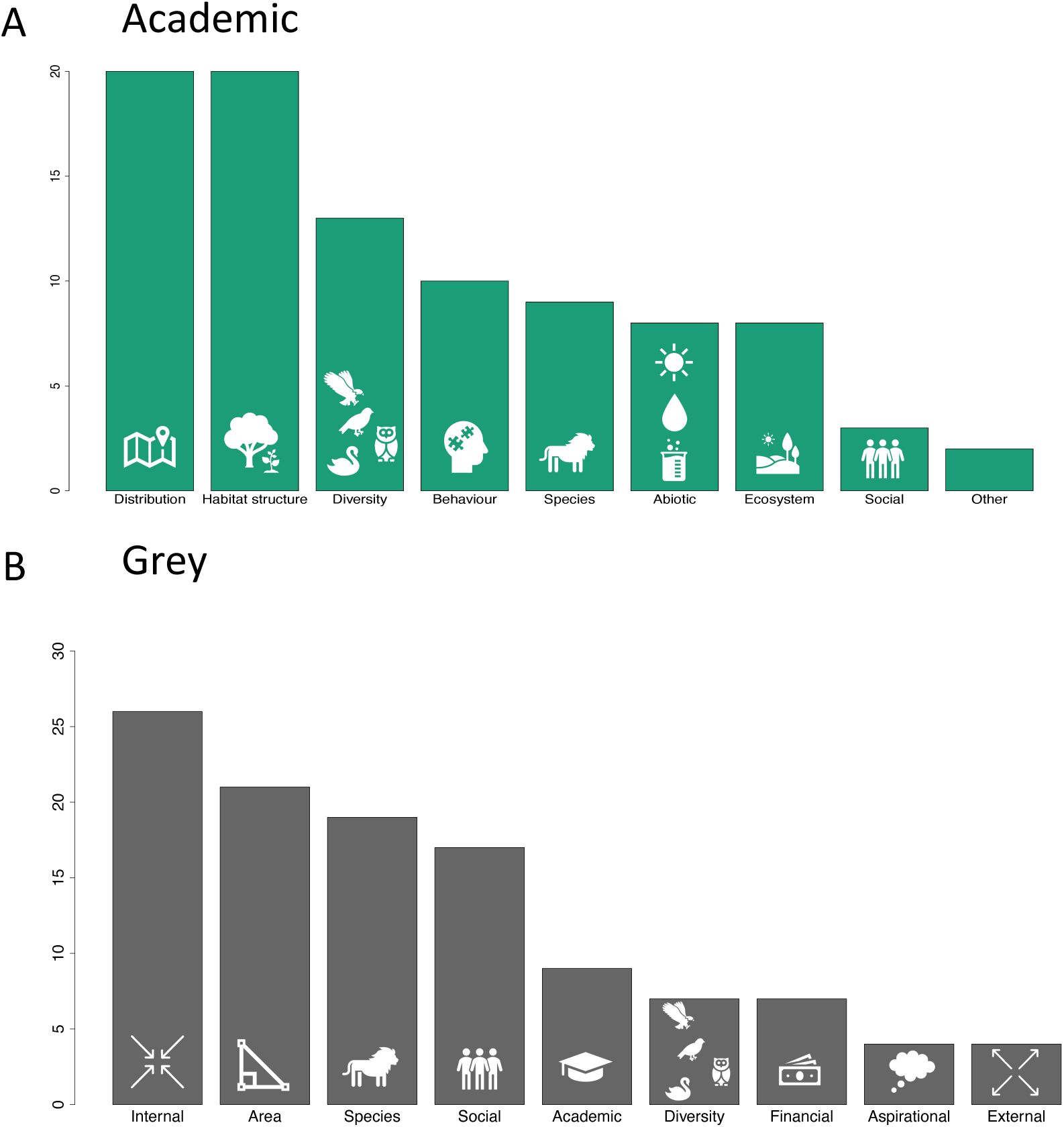
Differences between the response variables used to measure impacts of rewilding in academic literature and metrics used by rewilding organisations to evaluate progress. **A.** Response variables were extracted and categorised from vertical plot axes, figure and table legends, and / or sub-headings of results sections in academic case studies. Most metrics related to distribution (the location, distribution, density and / or abundance of a population) and habitat structure (including measures of vegetation independent of species composition, and measures of habitat taken through remote imagery, e.g. NDVI). **B.** Data was extracted from direct communication or analysis of grey literature. The metrics used most often to evaluate success in rewilding organisations are internal metrics (relevant within the organisation) and measures of land area. Measures related to species survival and population health, social outcomes, and diversity were common to both the academic and grey literature. Full detail on each category and how these were assigned can be found in supplementary information 1.

## Discussion

How monitoring and evaluation are conducted in rewilding and what can be concluded thus far is a ‘known unknown’ in the academic literature (Hart et al., 2023, Mahajan et al., 2023). Here we have mapped out the approaches to monitoring and evaluation in rewilding across both the academic and grey literature. We highlight some weaknesses and make recommendations for how they may be addressed. Broadly, this map suggests that there is some overlap between academic research and practitioners in terms of the methods used to collect data to monitor and evaluate rewilding, but we found divergence between how the two groups frame success and use evaluation metrics to measure rewilding’s impacts. This suggests that researchers and practitioners between them cover a greater range of rewilding’s impacts than is captured by assessment of each group separately, which demonstrates a strength of our approach. The divergence between researchers and practitioners is likely underpinned by the difference in contexts of the two groups -rewilding projects focus being driven by funding, policy and community factors, while research is more centred around novelty -as well as the constant challenge that measures of biodiversity are complex, multidimensional and difficult to quantify.

Because rewilding is not tied to an expected end-state in the same way as traditional restoration strategies (Durant et al., 2019; Pettorelli and Bullock, 2023; du Toit and Pettorelli, 2019) it has been described as ‘fluid and unscripted’ (Law, 2017) by proponents, and ‘vague’ and ‘plastic’ by critics (Jørgensen, 2015). As discussed, some consensus has emerged recently regarding the definition of rewilding (Carver et al, 2025a), but similar clarity has not formed around just what rewilding’s specific outcomes should be. This lack of clarity was reflected here in the disparity between the ways in which rewilding organisations frame success and the main foci of academic research (Figure 2), and in the vast range of metrics applied to measuring outputs for evaluation. Some form of consensus regarding what rewilding aims to achieve in *measurable* terms is now required to develop monitoring and evaluation strategies to form a reliable evidence base for rewilding. We do not presume to offer a firm definition for what success should look like in rewilding here, but we propose three ways in which future work might be designed to address current gaps in monitoring and evaluation in rewilding by maximising the advantages of the approaches already being taken by researchers and practitioners: (1) develop more comparative study designs, (2) standardise methods of data collection and recording, (3) work more collaboratively.

### 1. Develop more comparative study designs

Most academic case studies we reviewed were short-term and singlesite. We argue that research focus should move away from studying rewilding sites in isolation and instead prioritise comparison of multiple sites and / or intervention types, ideally covering longer time periods. Although no two sites will ever be identical in all aspects of site history, known differences in context (e.g. age, location, treatment, previous land use etc.) would be described and results presented in context, (following, for example, principles outlined for the design of natural experiments; McGarigal and Cushman (2002), Watts et al (2016). Comparative studies of this type would provide stronger insight into the generalisability of outcomes, adding greater value to understanding rewilding than simply increasing the number of isolated case studies on a few, well-described locations. A move towards greater use of experimental study designs would also allow quantitative evaluation as well as comparison (Ockendon et al., 2021).

Few rewilding projects are set up to include control landscapes, since scientific research is not their objective and establishing and managing experimental set ups would demand additional resources. Although it may be good practice to include control sites and BACI (before-after-control-impact) design in monitoring frameworks (Caruana et al., 2024), practitioners may resist investing time and money in control or alternative strategies if they are seeking to maximise the benefit of rewilding across their sites. From a research perspective, where rewilding projects are not set up with experimental design on site, studies could aim to include control systems where possible -for example a neighbouring, non-rewilded area if access is permitted. All these issues could be resolved with more money and land for conservation, but real-world, near-term solutions will require up-front recognition and reward for practitioners making use of evidence-based approaches to mitigate the impact such approaches may have on stakeholder needs and perceptions. Similar issues have been successfully solved in evidence-based conservation (Sutherland et al., 2019), and so similar approaches could work here.

### 2. Standardise methods of data collection and recording

In addition to designing studies that involve greater comparison, monitoring and evaluation methods should aim for increased standardisation to make findings easier to pool and compare. This can be achieved through more standardised collection methods and better recording and reporting of results.

We show here that fieldwork – direct, on the ground observation – accounts for most monitoring and evaluation in both the academic literature and in rewilding organisations (Figure 3). If researchers and practitioners can develop a common toolkit for data collection then larger datasets can be built to the benefit of all. Data collection methods should be simple, accessible, and scalable to allow comparison across broad and local scales (Levin, 1992), and methods will likely achieve best uptake if they are co-developed among researchers and practitioners, and built by consensus. There are methods currently in use that have been field-trialled in rewilding systems and published in the academic literature (*e.g.* Pates et al, 2025) that could be a starting point for developing and sharing good practice. Another approach may be to push for the widespread adoption of protocols currently in use by practitioners such as the British Trust for Ornithology’s breeding bird survey methods (BTO, 2025) and Australia’s Terrestrial Ecosystem Research Network protocols (TERN, 2025). Such methods are publicly available and already contribute to the development of a repository of biodiversity data (e.g. the BTO methodology is shared among the BTO, JNCC, and RSPB).

Improved data standards and storage would also allow maximum benefit to all stakeholders. In collating information for this systematic map from the academic literature, it was difficult to determine the exact location, age, and area of each case study described. Ideally, studies would include such details as standard, as well as information on the kind of rewilding intervention taken (methods, timelines, species reintroduced etc.), and the aims of the project from which the case study is drawn. In addition, continued progress towards open sourcing of information in both the academic (Mayo et al, 2016) and grey literature (Cadotte et al., 2025), and open access publication of research articles, will improve the way that information is stored and shared. Such steps would allow the development of repositories of good quality, comparable data on rewilding to allow conclusions to be drawn to inform future directions (Sutherland et al., 2020). This is particularly important in the context of our findings that the foci of researchers and practitioners are capturing different aspects of rewilding progress. Practitioners without easy access to developments in research are likely to share knowledge only through their own networks, meaning that valuable insights may be lost to both groups (Sabo et al., 2024).

### 3. Work more collaboratively

Of the rewilding organisations included in this review, most have objectives or definitions of success that they are aiming for (86%), most are collecting monitoring data (64%) and about half are evaluating progress (52%). The gap between monitoring and evaluation suggests that more information may be being collected than is being used to inform action. As well as improved data recording and sharing (including open sourcing of data), better collaborative relationships between researchers and practitioners could help to put more information to use in building the largest possible picture of what is working and where in rewilding. When beginning a relationship with a rewilding organisation to arrange study site access, researchers could request and review historical data relevant to their study aims. Existing findings can then either be folded into a study if appropriate or acknowledged as background, recognising that data and findings are valuable and it is important that practitioners also benefit from data sharing (e.g. data sovereignty where appropriate). Development of such relationships could serve to incentivise positive practitioner-researcher collaborations such as establishment of long-term ecological plots for study.

A step that rewilding organisations could take where possible is the development of SMART targets to direct monitoring efforts and support development of metrics to measure progress (Aldridge and Colvin, 2024). This would also provide a scaffold for academic researchers to consider when designing studies of their sites. Where possible, research could be aligned with - or at least set within the context of -the aims of the rewilding project in which they are situated. Researchers building study proposals jointly with practitioners would allow both groups to design work targeted to gaps in the knowledge base. Our results suggest that this may involve the inclusion of more discussion of social considerations in academic research as social and economic factors are currently receiving less attention than ecological factors, but the former often drive the work of rewilding projects (Figure 2). Taking steps to improve collaboration and share data between researchers and rewilding organisations would develop relationships while adding depth and context to the academic literature.

## Conclusions

There is strong academic and practitioner interest, as well as international political will, to address the damage done to ecosystems worldwide (FAO, IUCN, CEM and SER, 2021). This is good news for conservation, but funding and other resources will – rightly – be directed towards strategies that can be proven effective. In particular, robust data will be needed to reassure financial mechanisms for securing long term investment in rewilding strategies (Rewilding Britain, 2024), and the development of a strong evidence base for rewilding will allow practitioners to ensure that effort and resources are applied effectively (O’Connell and Prudhomme, 2024). Monitoring and evaluation are key here (Hart et al., 2023, Svenning et al, 2016). To meet the scale of the challenge of the environmental crisis, researchers and rewilding practitioners need to know what works and where. Good practice can only be established and shared responsibly if it is based on evidence (Segan et al., 2011, Sutherland et al., 2020), so the challenge is now to establish the evidence base sufficiently to direct the development of future action. That means accruing more comparable data from a broader range of contexts, and we have outlined some possible avenues to begin.

We acknowledge that many of our findings reflect known problems with monitoring and evaluation not just in rewilding, but more broadly in restoration and conservation. There have been calls for improved study design and use of controls in ecological research for many years (Sutherland, 2022, Christie et al., 2019) and similar problems have been raised for practitioners (Legge, 2015). The improvements we suggest are not simple to achieve, especially for rewilding given the large spatial and long temporal scales involved. Control data for academic studies will likely need to come from landowners other than those of the rewilding sites, requiring additional permissions to survey wider areas, and this will add complexity to planning processes. Furthermore, realities of funding and resources in academia tend to limit data collection. The large number of studies we found here that cover only one or two years of data likely results from short-term scientific funding cycles. Alongside our call for more comparative studies, improved data collection and recording, and more collaborative relationships between researchers and practitioners, we call for greater funding. In particular, projects that extend the spatial and temporal reach of monitoring and evaluation should be championed. If, as a research community, we can start to implement the approaches outlined here, we will begin to build a body of academic literature that can tell us where and how rewilding is working – achieving what we want (and need) it to.

## Supporting information

Supplementary 2 - extended results

## Acknowledgements

WDP and the Pearse Lab are funded by UKRI BB/Y008766/1, NE/X00547X/1, NE/X013022/1, the Alan Turing Institute, the Singapore Green Finance Centre, and Hitachi Ltd. NJ Pates is funded by the Imperial President’s PhD scholarship. JMB was funded by the UKCEH National Capability for UK Challenges programme NE/Y006208/1. NP is funded by Research England. We would like to gratefully acknowledge the assistance of Jingyi Yang, Cecilia Maria Montauban and Jonas Lembrechts in translating portions of literature for inclusion in this study. We are also grateful for the assistance of all the Global Rewilding Alliance partner organisations who provided their time to contribute information to this study.

