## Supplementary 1 - extended methods for "Systematic mapping shows monitoring, evaluation, and engagement are needed to build an evidence base for rewilding"

### Supplementary information 1

This supplementary information contains extended methods for: **Systematic mapping shows monitoring, evaluation, and engagement are needed to build an evidence base for rewilding**

James M. Bullock [b],

Johan T. du Toit [c],

Sahran Higgins [d],

Nathalie Pettorelli [b],

William D. Pearse [a],

#### Extended Methods

This section of supplementary information explains how decisions were made to classify academic studies and information provided by rewilding organisations into categories for analysis in this study.

We would like to be clear that the classification of a study or organisation into any of the groups outlined below is in no way an objective statement on the quality of any study, initiative or organisation. Each study and organisation included has its own context, aims and objectives. These do not necessarily align fully with the specific research questions we set out to answer. Classifications, therefore, reflect the best assessment possible using the criteria below and the information available at the time of writing.

### 1. Academic literature

#### 1.1 Screen for academic literature

Academic literature was screened in multiple stages (Table S1.1). We downloaded titles and abstracts of all search results, pooled them, and removed duplicates. Titles and abstracts were screened using the R package metagear (Lajeunesse, 2016) using pre-determined criteria. We aimed to conserve as much data as possible at each stage, so a 'maybe' category was used in addition to 'yes' and 'no' to carry uncertain cases forward to a full review.

To be included at the first stage, articles had to meet two criteria:

(i) Be a piece of primary research or a meta-analysis. Opinion or perspective pieces, for example, were excluded.

(ii) Include a description of a rewilding case study. Most literature reviews were therefore excluded, as was research that used computer modelling alone to predict future rewilding outcomes. One review paper, Thomas et al. (2023), was included because it compiled and analysed the outcomes of multiple rewilding case studies.

The final stage of screening was full paper review, where each paper was compared against a further set of criteria for inclusion:

(i) Confirm a piece of primary research or a meta-analysis.

(ii) Confirm contains a description of a rewilding case study.

(iii) Contains detail on either monitoring, or evaluation, or both, of the rewilding case study (in line with our definitions).

One thesis chapter was included following the screening process (Beason, 2020). Another relevant thesis linked to two additional papers that were not found in the initial screen and these were added to the total (Warner et al., 2021, Warner et al., 2022).

*Cases requiring additional consideration:*

We required papers to explicitly describe their studies as rewilding as this is the key focus of this study. One borderline paper did not mention rewilding by name (referring instead to a 'large-scale refaunation effort') however, 'rewilding' was one of the article's keywords and it was therefore included (Correia et al., 2017). A second borderline paper was excluded from the data set as it stated plainly that is was 'not a rewilding scenario' (Hall and Bunce, 2019).

Three papers were added to the data set while conducting the grey literature review as rewilding organisations provided academic papers in support of their monitoring and / or evaluation activity. To collate the largest possible selection of data, these were included provided they conformed with the criteria above *except for the requirement of explicit mention of rewilding*. These were case studies on forest restoration and species reintroductions (Barri et al., 2021, Becerra et al., 2019, Walker et al., 2022). Although these papers did not reference rewilding, since they were provided as evidence by rewilding organisations involved in the content of those studies, they were deemed appropriate for inclusion.

**Table S1.1 A total of 48 papers were included in the academic literature review**. The results of an initial search and subsequent screening of papers discovered in four academic databases using search string "rewild* AND (evaluat* OR monitor*)". All papers found by the initial search were downloaded and duplicates removed. Two rounds of screening followed: first by title and abstract, then by full paper review. Two papers were found in a PhD thesis that were added to the total as they met the criteria specified. Three more were contributed by members of the Global Rewilding Alliance who were contacted as part of the grey literature review.

| | **Initial search**<br>**n =** | **Remove duplicates**<br>**n =** |
|---|---|---|
| **Web of Science** | 265 | 265 |
| **Scopus** | 221 | 52 |
| **PubMed** | 79 | 9 |

| | | |
|---|---|---|
| **Open Access Theses and Dissertations** | 16 | 14 |
| | TOTAL | 340 |
| Title and abstract screen | | 96 |
| Full paper review | | 43 |
| **Total** including additional literature found through other means | | **48** |

### 1.2 Extract for academic literature

Table S1.2 explains how each information type was extracted from papers and coded for the academic literature review. Direct extraction was the preferred option where possible. In classifying studies as ‘monitoring’, ‘evaluation’, or both, these categories were assigned based on our definitions, regardless of how studies framed themselves within the text. Additional information on extraction and coding, and some illustrative examples are provided below.

#### 1.2.1 Country

The country listed is the country of the case study described in the article. In cases where an article described multiple case studies in different countries, e.g. Segar et al. (2022), each country is recorded separately.

#### 1.2.2 Location

If provided, the precise location of a site / sites was extracted. These were recorded as provided and were also converted into latitude and longitude to allow mapping. Where multiple precise locations were provided, each was recorded separately, e.g. Kaštovská et al. (2024). Where precise locations were not provided, e.g. Thomas et al. (2023), coordinates were recorded as the approximate centre of the country of the case study / studies to allow mapping.

#### 1.2.3 Study area and Rewilding area

All areas and area estimates provided within articles were extracted and then interpreted according to the rest of the information available within the text and figures. We converted all areas to square kilometres from other units as required. In some cases, authors were clear in describing their study locations (e.g Li et al. (2018) specified a study area of 45$km^2$ within a much larger reserve of 152,300$km^2$). However, where only a single area was provided, it was not always possible to determine whether this referred to the area studied, or simply to the size of the rewilding project within which the study was situated.

To account for this, we recorded ‘study area’ and ‘rewilding area’ where a distinction could be made. If only one area was provided but there was sufficient context then we assigned it to either the broader rewilding area, or a narrower study area. We were primarily interested in the area investigated in the case studies described, so additional columns ‘area certain’ and ‘area uncertain’ were added. **Area certain** contained the areas provided where we could be reasonably sure that they referred to the area studied in the reported case study and this number was used in analyses.

#### 1.2.4 Monitoring or evaluation

Some articles defined themselves as monitoring, some as evaluation, some as both, and some as neither. This category was assigned according to the definitions established during the scoping of this study:

- ***Monitoring*** *is the practice of gathering data over time. To be classified as a monitoring study here, the paper must make repeated, comparable measures over time. Usually, this means repeated field seasons.*
- ***Evaluation*** *is the process of assessing whether an action – in this case, rewilding – has achieved the desired outcome. To be classified as an evaluation study here, therefore, the paper must make clear the aims of the intervention and whether the results of the study were in-keeping with those aims.*

<u>Illustrative examples:</u>

- Merelli et al. (2024) describe their paper as both monitoring and evaluation. However, all data was collected in a single fieldwork season and time was not incorporated into their analyses. It does not, therefore, meet the definition of monitoring used here and it was categorised as evaluation only.
- Li et al. (2018) do not mention monitoring in their paper. However, as their data set comprises breeding-season bird surveys repeated over 2 years, it is classified as both monitoring and evaluation here.

#### 1.2.5 Interest

This category was assigned to capture the main study system of interest in each case study (plants, mammals, birds etc.). 9 categories were used: mammals, plants, birds, invertebrates, reptiles, social, abiotic, ecosystem, and fungi. Categories were assigned based on interpretation of abstracts and introductions, as well as the focus of response variables.

The 'social' category involved people (treated as a separate category from mammals). These studies tended to focus on cultural or social aspects of rewilding, e.g. Heuer et al. (2023), Quintas-Soriano et al. (2023). 'Mammal' studies, on the other hand, were more often focused on population responses such as distribution (Hurtado et al., 2020, Mata et al., 2021) and survival probability (Smith et al., 2023, Thomas et al., 2023).

Although most studies discussed the restoration of whole ecosystems, only two were categorised as 'ecosystem' for their main interest (Segar et al., 2022, Torres et al., 2018). These two studies conducted interviews and questionnaires to base findings on a range of factors contributing to overall scores for 'ecological integrity' and 'human forcing' and could not be easily separated into one of the smaller categories of interest.

Most studies were given a single category, however, it was sometimes appropriate to assign more.

<u>Illustrative examples:</u>

- *Reptiles and plants.* Tapia and Gibbs (2023) describe a rewilding project with giant tortoises where the main response variables were related to tortoise distribution and vegetation responses.

- *Mammals and abiotic*. Kaštovská et al. (2024) focus mainly on abiotic responses (soil dry weight, pH, soil carbon, nitrogen, and phosphorous stocks etc) in their study, but their results are framed by a project rewilding with ponies, bison and cattle.

- *Plants, invertebrates, birds*. Warner et al. (2021) describe the responses of three groups to rewilding by reforestation: plants, carabid beetles, and birds.

**Table S1.2 A key to the process followed to extract and code information from academic papers to inform the systematic map.** Three categories of information were extracted: bibliographic information (green highlight); case study information (blue highlight); research question information (orange highlight). Rows are the column headings in the full data set. Columns describe how data was extracted and / or interpreted to allow collation and comparison. Direct extraction means data could usually be copied straight from the article without any requirement for interpretation (all bibliographic information, for example). Later columns explain the process followed where interpretation was required to assign classifications through coding.

| | Direct extraction | Or… | Then |
|---|---|---|---|
| ***year*** | *Bibliographic info* | | |
| ***title*** | *Bibliographic info* | | |
| ***authors*** | *Bibliographic info* | | |
| ***country*** | *Title / abstract / intro / methods* | | |
| ***location*** | *Title / abstract / intro / methods / figure legend* | *Central location on country chosen from Google Maps* | |
| ***study area*** | *abstract / intro / methods / figure legend* | *Interpreted from areas provided and context (methods, figure legend)* | |
| ***rewilding area*** | *abstract / intro / methods / figure legend* | *Interpreted from areas provided and context (methods, figure legend)* | |
| ***monitoring or evaluation*** | *Title / abstract / into / discussion / conclusions* | *Interpreted from areas provided and context (methods, figure legend). Direct extraction was not the primary method of classification here, see additional information and illustrative examples.* | |
| ***keywords*** | *Keywords* | | |
| ***interest*** | | *Interpreted from title / keywords / abstract / results* | *Grouped into 9 broad categories: mammals, plants, birds, invertebrates, reptiles, social, abiotic, ecosystem, fungi* |
| ***control*** | *Methods* | *Interpretation from methods / figure legends* | |
| ***sites*** | *Methods* | *Interpretation if necessary* | |

| | | | |
|---|---|---|---|
| ***duration*** | *Abstract / introduction / methods* | *Inferred if not explicit* | |
| ***methods*** | *Methods* | | *Grouped into 7 broad categories: fieldwork, remote sensing, wildlife tracking, experiment, consultation, DNA methods, review* |
| ***response variables*** | *Methods / results / figure legends* | | *Grouped into 9 broad categories: distribution, habitat structure, diversity, behaviour, species, abiotic, ecosystem, social, other.* |

#### 1.2.6 Control

To understand how confident we can be in assessing the relationship between rewilding interventions and the results described, the control column recorded YES or NO. To be coded YES, studies needed to meet one of two criteria:

(i) Full description of a study design / treatment(s) in which an experimental control was described, e.g. Tapia and Gibbs (2023), Temmink et al. (2022).

(ii) Sufficient detail in methods to establish that results compare different treatment types, one of which represents a version of ‘no change’.

<u>Illustrative examples:</u>

- *YES*. Schulte to Bühne et al. (2022) included the land around the rewilding site in their study to compare changes within the rewilded area to those outside. Thers et al. (2019) compared results in spontaneously regenerating forests with those in planted and managed forests.
- *NO.* Dombrovski et al. (2022) consider changes in bird species composition within a rewilded area, but do not include data from outside. Beason (2020) describes 3 habitat types in their study, but as all are under rewilding management, this does not constitute use of a control by our definition.

#### 1.2.7 Sites

We recorded the number of rewilding case studies included in articles to understand how often multiple rewilding sites are compared and contrasted. This was typically clearly stated in the methods sections of articles (e.g. Segar et al. (2022) describe 7 case studies).

#### 1.2.8 Duration

To understand the length of time spent studying rewilding effects in academic studies, we extracted duration. Categories represent number of years in whole numbers. In most instances, this was simply extracted from methods as the duration of sampling or fieldwork that went into collecting the data. There were some cases where duration was not simple to assign, and the following criteria were used:

(i) Where a study compared snapshots, we counted the time spanned by the window.

Illustrative examples:

- Some studies using remote sensing data compared an early snapshot with the present. E.g. Schulte to Bühne et al. (2022) studied vegetation cover over 20 years (duration = 20 years). E.g. Rincon-Madroñero et al. (2024) used 30 years of NVDI data (duration = 30 years).
- Konvička et al. (2021) used presence-absence data from 3 time points, 1994, 2009, and 2016-19. The span from the earliest of these dates (1994) to the latest (2019) was 25 years.
- In some cases, although a window of time was described, the data included did not support the use of the timespan for duration. Segar et al. (2022) describe case studies where rewilding has been happening for several years (between 2011 and 2014-2020), however, they elicited only expert comment in the year of the study *about* the baseline years. This study was categorised duration = 1. Similarly, Thomas et al. (2023) presented a review of case studies conducted between 2007 and 2021. Their published work, though, represents a single review, duration = one year.

(ii) Where a single study reported data collected over different lengths of time, the longest duration was recorded. E.g. Tapia and Gibbs (2023) collected vegetation mapping data over 15 years, and eight years of plant community composition data (duration = 15 years).

#### 1.2.9 Methods

Methods were extracted directly from methods sections into one column (methods named), then grouped by coding into one of 7 categories: fieldwork, remote sensing, wildlife tracking, experiment, consultation, DNA methods, review (Table S1.3). This allowed us to draw conclusions about the way in which studies are conducted. More than one method category was assigned where appropriate.

Illustrative examples:

- Methods described in Smith et al. (2023) included experimental trials (experiment), use of live relay cameras (wildlife tracking), species recapture (fieldwork), use of remote microchip scanners (wildlife tracking), and radio tracking (wildlife tracking). This study was coded fieldwork, experiment, wildlife tracking.

**Table S1.3 Methods used to monitor and / or evaluate rewilding case studies in the academic literature were coded according to 7 broad categories: fieldwork, remote sensing, wildlife tracking, experiment, consultation, DNA methods, review.** The methods described (extracted directly from papers into the column 'methods named') in each study are shown here (duplicates removed), assigned to the relevant code.

| Fieldwork | Consultation | Remote sensing | Wildlife tracking | Experiment | DNA methods | Review |
|---|---|---|---|---|---|---|
| Breeding bird surveys | Delphi technique | Satellite remote sensing | GPS wildlife tracking | Herbivore exclosure | eDNA metabarcoding | Literature review |
| Vegetation measures | Questionnaires | Satellite imagery | Radio telemetry | Experimental trials | DNA barcoding | |
| Censuses (counting) | Interviews | Land cover classification | Camera traps | Full-factorial field experiment | | |
| On ground observations of animals | Workshops | Primary productivity analyses | Radio tracking | Transplant experiment | | |
| Sample collection (sticks, dung…) | Feedback solicitation | Lidar | Mark-resighting | Tea bag decomposition | | |
| Transect surveys | Cultural monitoring survey | Habitat categorization | Remote camera observation | | | |
| Habitat surveys | | NVDI | Animal capture | | | |
| Habitat mapping (UAV) | | Habitat mapping (UAV) | Live relay camera | | | |
| Mark-resighting | | | Remote microchip scanners | | | |
| Plant community surveys | | | Radio tracking | | | |
| Pitfall traps | | | Passive acoustic monitoring | | | |

| |
|---|
| Earthworm counts |
| Animal capture |
| Moth light traps |
| Driving transects |

#### 1.2.10 Response variables

The same approach to the analysis of methods was followed for analysis of response variables. Named response variables were extracted directly from plot y-axes, figure and table legends, and / or sub-headings of results sections. These were then grouped into 9 broad categories for analysis: distribution, habitat structure, diversity, behaviour, species, abiotic, ecosystem, social, other (Table S1.4). More than one category was assigned where appropriate. Greater detail on the coding assigned is provided below:

**Distribution.** Response variables related to the location, distribution, density and / or abundance of a population. This category includes measures used as proxies for the same, for example, abundance of faeces in exclosure experiments (Tapia and Gibbs, 2023). 'Spatial distribution of open-land habitat' and 'spatial distribution of tree species' were borderline cases, both from the same source (Zielke et al., 2019). While these responses could also have been coded as measures of habitat structure, as they were broken down in figures to species level they were deemed closer measures of species distribution and density and so were captured in the distribution category.

**Habitat structure**. Response variables related to the structure of the system. This category includes measures of vegetation that are independent of species make-up, and measures of habitat taken through remote imagery, e.g. NDVI.

**Diversity**. Measures and metrics related to species diversity and community composition.

**Behaviour**. Measures informed by direct observation, or observation-based modelling, of species or population behaviours.

**Species.** Responses related to individuals within a focal species. These were commonly found in studies of species reintroductions. Example responses include number of translocated individuals, percentage of individual animals released using different protocols (Thomas et al., 2023), individual body masses of released animals (Smith et al., 2023) etc. Context informed some classifications. For example, measures of prey composition, prey preference, and habitat use could fall under diversity or behaviour categories, but because they were listed as the results of a cheetah release study, they were recorded under the species category (Marker et al., 2024).

**Abiotic.** Responses concerned with physical, rather than biological responses, e.g. measures of light, temperature, physicochemical soil aspects, etc.

**Ecosystem**. Measures combining multiple aspects of a system or its function were included in this category, for example: rewilding score, ecological integrity, and human forcing (Segar et al., 2022, Torres et al., 2018). Also included were measures of an ecological function e.g. decomposition (Warner et al., 2022) or ecosystem respiration (Temmink et al., 2022), or measures investigating species or ecological interactions within a system (Fernandez et al., 2017).

**Social.** Measures involving the human response to rewilding case studies were listed as ‘social’. These typically involved assessments of societal perceptions of rewilding interventions e.g. Quintas-Soriano et al. (2023), Pettersson and de Carvalho (2021).

**Other.** The few measures (n = 4) that did not fit any of the above categories were placed into the ‘other’ category. Two measures came from a study investigating rewilding by studying vertebrate visitors to communal rhinoceros latrines; in addition to diversity and behavioural measures, the study reported on latrine quality and size, neither of which fit sensibly into other categories (Awasthi et al., 2024). The other two responses were the number of reintroduction projects and the objectives of those projects (Banasiak et al., 2021).

**Table S1.4 Response variables extracted from rewilding case studies in the academic literature were coded into 9 broad categories: abiotic, habitat structure, distribution, species, diversity, behaviour, ecosystem, social, and other.** The variables as extracted directly from papers in each study are shown here (duplicates removed), assigned to the relevant code.

| Abiotic | Habitat structure | Distribution | Species | Diversity | Behaviour | Ecosystem | Social | Other |
|---|---|---|---|---|---|---|---|---|
| Soil moisture | Land cover class accuracy | Density of breeding raptors | Number of translocated individuals | Raptor community change | Predicted habitat selection | Rewilding score | Local stakeholder perceptions | Number of reintroduction projects |
| Light | % area cover | Change in raptor breeding pair abundance | % individual success rate (survival > six months) | Realized and potential faunal complexity score | % Behaviours | Ecological integrity | Rewilding progress and threat factors | Reintroduction objectives |
| Nitrogen | % area cover change | Raptor composition by habitat type | % captive-born, period of captivity, and wild-born | Species ID | Daily activity pattern | Human forcing | Socio-cultural characteristics | Latrine quality |
| CO2 flux | Normalized Difference Vegetation Index (NDVI) | Correlation plot of raptor species | % hard- and soft-release | Species richness | Food selection | Ecological interactions | Perceptions of landscape changes | Latrine size |
| CH4 flux | Change in vegetation dynamic parameters | Realized and potential occupancy of reintroduced species | Effects of fence / origin / release / sex / age | Endemic diversity | % habitat use for food selection | Species interactions | Perceived drivers of land use change | |
| Soil dry weight | Land cover change | Area and proportion of realized and potential occupancy | Body mass | Avian diversity | Potential habitat preferences | Network of seed dispersal | Perceptions of nature's contributions to people | |
| Soil bulk density | Habitat classification | Tortoise feces | Population survival probability | Avian assemblage | Bison movement | Animal linkage level | | |
| % clay | Grass cover | Cactus cladodes | Probability of initial occupancy | Community composition | Bat activity | Net ecosystem exchange | | |
| pH | Forb cover | Cactus fruits | Probability of colonisation | Species accumulation | Bat and bird activity by habitat type | Ecosystem respiration | | |
| Index WSA | Woody plant regeneration | Detection probability | Frequency of detection | Plant count | Bat passes per night | Net nitrification rate | | |

| | | | | | | |
|---|---|---|---|---|---|---|
| Soil water retention capacity | PCA values for lidar-derived cell values | Occupancy probability | Weight | Shannon diversity | Behaviour rate | Respiration rate |
| Soil C stock | Trend in NDVI | Density | Apparent survival | Acoustic complexity index | Rate of establishment | Decomposition |
| Soil N stock | Habitat distribution | Occupancy vs density | Growth rate | Bioacoustic index | Single-season site occupancy | Soil invertebrate feeding |
| Soil P stock | Landscape heterogeneity | Spatial distribution of tree species | Release success | Specialist richness | % time vigilant | |
| Soil C/N/P ratios | Changes in environmental composition | Spatial distribution of openland habitat | Time to independence | Rank-abundance | Site fidelity | |
| Soluble C | Changes in vegetation parameters | Species abundance | Prey composition | Simpson index | | |
| Soluble N | Woody plant regeneration | Endemic abundance | Prey preference | Absolute effective diversity | | |
| Soluble reactive P | Herbaceous plant cover | Change in occurrence rate of bird species | Hunting success | Composition of latrine visitors | | |
| Microbial C | Opuntia fruit, cladode, and regeneration | Population numbers | Habitat use | Genetic diversity | | |
| Microbial N | Forest cover | Herd demographics | Step length | Spore morphospecies | | |
| Microbial C/N ratio | Vegetation cover | Population size | Daily distance travelled | Species traits | | |
| Portion of fine particles | Reed shoot number | Population density | Speed | | | |
| Grain size | Maximum reed height | Mean coverage by species | Distance | | | |
| Dry bulk density | Vegetation biomass | Dispersal distance | Survival | | | |
| Organic matter | Shoot height | Number of cacti | Reproductive success | | | |

| | | |
|---|---|---|
| Carbon content | Shoot density | Iguana and tortoise feces |
| Sediment deposition patterns | Plant coverage | Plant density |
| Aboveground tree carbon | Distance to water | Prey density |
| Topsoil carbon | Seedling regeneration | Frequency of latrine visitors |
| Topsoil nitrogen | Ground layer height | Home range size |
| Nutrient concentration | Moss depth | Home range |
| % Bare soil | Tree biomass | Residence area size |
| Total C | Anthills | Population trends |
| Total N | Grass height | Spore density |
| C:N | Lagomorph droppings | |
| Soil texture | Lagomorph excavations | |
| Electrical conductivity | % AMF root colonisation | |
| Extractable P | | |
| Nitrates | | |
| Gravel | | |

### 2. Grey literature

#### 2.1 Collect for grey literature

We collected data from the screened subset of Global Rewilding Alliance (GRA) partners in two stages.

##### 2.1.1 Direct contact

First, we contacted organisations directly with an email:

- Email 1: 16 October 2024

*I am a researcher at Imperial College London and I hope that you can help me with a review of how monitoring and evaluation is happening in rewilding projects. Please could you send me any documents you have on how you monitor and/or evaluate the rewilding success of your projects, or put me in touch with someone at your organisation who can?*

*I feel that organisations like yours, that are engaged in the stewardship and conservation of land, must have a lot of information on this subject, and I want to bring as much of it together as possible to review practice and progress on a large scale. I am contacting all partner organisations in the Global Rewilding Alliance that have projects involving landscape management. I would be very grateful for anything that you could provide to help me assess:*

- *How your organisation defines success in rewilding. Is there anything specific that you have set out to achieve and how did you choose this?*
- *How you monitor progress. What do you record, and how often? What do you do with that data?*
- *How you evaluate your project. In the context of #1 and #2 – can you say how far along you are towards your definition of success? And how do your monitoring efforts feed into your evaluation?*

*All data used will be anonymised (I am happy to acknowledge your contributions if you prefer but will default to anonymity unless I hear otherwise from you).*

*Thank you very much for your time,*

- Email 2: 7 November 2024

*This is a follow up email to ask for help with a review of how monitoring and evaluation is happening in rewilding projects. I am a researcher at Imperial College London and I hope that you can send me any documents you have on how you monitor and/or evaluate the rewilding success of your projects, or put me in touch with someone at your organisation who can?*

*I want to bring together as much information on this topic as possible to review practice and progress on a large scale. I am contacting all partner organisations in the Global Rewilding Alliance that have projects involving landscape management. I would be very grateful for anything that you could provide to help me assess:*

- *How your organisation defines success in rewilding. Is there anything specific that you have set out to achieve and how did you choose this?*
- *How you monitor progress. What do you record, and how often? What do you do with that data?*
- *How you evaluate your project. In the context of #1 and #2 – can you say how far along you are towards your definition of success? And how do your monitoring efforts feed into your evaluation?*

*All data used will be anonymised (I am happy to acknowledge your contributions if you prefer but will default to anonymity unless I hear otherwise from you).*

*Thank you very much for your time,*

#### 2.1.2 Documents

Secondly, we collected publicly available documents, either provided by respondents to the email approach, or through websites. Websites were searched for documents – annual reports, progress reports, impact statements. If reports for multiple years were available, the most recent was used. Documents were searched using the table of contents, as well as searching within for ‘monitor’, ‘evaluate’, ‘measure’, ‘track’, ‘impact’, and ‘assess’.

Where documents were not available, information was extracted directly from webpages. Key search areas were home pages, ‘About us’, ‘Our work’, ‘projects’, ‘our impact’ etc.

### 2.2 Extract for grey literature

Table S1.5 explains how information was extracted from emails, notes from conversations, and documents, and then coded for the grey literature review. Additional information on extraction and coding is provided below. Fewer illustrative examples are provided here and in analyses of the grey literature to limit identification of individual organisations and protect their privacy.

Most written information from GRA organisations was available in English, either because it was produced in English, or because websites provided multiple language documents, or had in-built translation functions. In these cases, the English language version was used for analysis. In 8 cases, it was not possible to access English language documents: three in Spanish language, three in Portuguese, one in Chinese, and one in Dutch. We sought native speakers of these languages and explained the protocol to them to extract data for these organisations. We were not able to find a native Portuguese speaker, so these three sites and their documents we used Google Translate to extract data.

#### 2.2.1 Area (km$^2$)

All areas provided or found within documentation were converted into square kilometres from other units as required. If multiple projects were described, their areas were combined. Areas were grouped into the same categories used for the academic literature.

#### 2.2.2 Start year and Duration

To understand the length of time different rewilding organisations have been operating, we took the start year listed in documentation and subtracted that from 2025 to give duration. If an organisation had been running or adding short-term projects over the span of its existence, we used the earliest year provided / the year of the organisation’s founding or establishment.

**Table S1.5 A key to the process followed to extract and code information from grey literature and responses to inform the systematic map.** Four categories of information were extracted: partner organisation detail (green highlight); direct communication detail (blue highlight); case study information (purple highlight); research question information (orange highlight). Rows are the column headings used in the full data set. Columns describe how data was extracted and / or interpreted. Direct extraction means data that was copied straight from an email response, website, or document without requirement for interpretation. Later columns describe the coding that was then assigned to allow collation and comparison.

| | Direct extraction | Or… | Then |
|---|---|---|---|
| ***Continent*** | *Global Rewilding Alliance (GRA) website* | | |
| ***Organisation*** | *GRA website* | | |
| ***Operating in*** | *GRA website* | | |
| ***Response*** | | *Y/N assigned depending on whether a response was received (beyond automatic reply) to either email* | |
| ***Provided*** | *Email content.*<br>*Link from correspondence.* | | |
| ***Document / page*** | *Attachment or link from correspondence.* | *AND / Or documents publicly available online. E.g. annual report, impact statement, or pages on websites e.g. 'About us', 'Impact', 'Our work' etc.* | *Information from both response if provided and documentation included in analysis.* |
| ***Area (ha)*** | *Correspondence, documentation, and / or conversation.* | *Consolidated if possible from separate project pages,* | |
| ***Start (year)*** | *Correspondence, documentation, and / or conversation.* | *Online search for 'When was [GRA Partner Name] established?'* | |
| ***Duration*** | | *Calculated as: 2025 – start year* | |
| ***Define success*** | | *YES assigned if a definition or set of objectives available* | |
| ***Success direct extract*** | *Correspondence, documentation, and / or conversation.* | | |

| | | | |
|---|---|---|---|
| ***Success classification*** | | | *Grouped into 8 broad categories: social, knowledge sharing, ecosystem, species, diversity, habitat structure, abiotic and other* |
| ***SMT*** | | *Combination or S, M, and T assigned for definitions of success that are smart, measurable, and / or time-bound.* | |
| ***Monitor*** | | *YES assigned if information on current monitoring available* | |
| ***Monitor direct extract*** | *Correspondence, documentation, and / or conversation.* | | |
| ***Monitor classification*** | | | *Grouped into 7 broad categories: fieldwork, remote sensing, wildlife tracking, DNA methods, consultation, experiment, and aspirational* |
| ***Evaluate*** | | *YES assigned if information on current evaluation available* | |
| ***Evaluate direct extract*** | *Correspondence, documentation, and / or conversation.* | | |
| ***Evaluate classification*** | | | *Grouped into 9 broad categories: internal standards, social, species, academic, diversity, financial, area, external standards, aspirational* |

#### 2.2.3 Define success

'YES' or 'NO' was recorded in this column to understand the number of organisations working towards a stated goal, objective, or definition of success.

#### 2.2.4 Success direct extract and categorisation

Definitions of success were taken directly from documents, websites, or direct communication with GRA organisations. Within documents, extraction was typically from stated organisational mission statements, goals or objectives. If these were not available, organisational 'vision' statements, or statements from introductory pages of annual reports sometimes provided a definition of success. This data was often long-form, detailed and not easy to compare across organisations. However, retaining it allows easier checking of classification and SMT categorisation so it was kept in the form of the direct extraction column of the data set.

Definitions were coded into eight broad categories: social, knowledge sharing, ecosystem, species, diversity, habitat structure, abiotic and other (Table S1.6). More than one category was assigned where appropriate. If responses via email / interview and the data provided online were different, then all data was combined and coded for analysis. Greater detail on the coding assigned is provided below and in Table S1.6.

**Social.** Definitions involving community and socio-economic impacts were classed as 'social'. This included definitions detailing ecosystem services (healthier air, water etc for communities); tourism, wellbeing, connection to nature, stewardship of the land, employment and other financial opportunities etc.

**Knowledge sharing.** Many organisations tied success to the creation and sharing of protocols, methods, research, data etc to allow others to learn from good practice and undertake rewilding.

**Ecosystem**. Many definitions of success referred to functioning, thriving ecosystems and / or processes. We also included success related to climate change mitigation and resilience in this category.

**Species.** Several organisations are built around threatened, reintroduced, or charismatic species and thus many definitions of success were related to the progress of those focal species. This category captures definitions that name species, or list measures of success related to survival, reproduction, establishment etc.

**Diversity**. Definitions of success related to species diversity and community composition.

**Habitat structure**. Measures of success related to area and connectivity of land managed by an organisation were classified under habitat structure.

**Abiotic.** One measure of success was concerned with physical, rather than biological success: reduction of emissions and increased carbon sequestration.

**Other.** Several definitions did not fit any of the above categories and were placed into the 'other' category. Some were definitions of success tied to financial progress or success *of the rewilding organisation* (many other considerations related to money were categorised under 'social' when they involved benefits to the wider community). Some were legal considerations, related to permitting, and / or securing legal recognition and protections for land. And some were measures of success related to the achievement of targets, which did not fit well into any of the other categories.

#### 2.2.5 SMT

Because it is difficult to compare across rich, narrative data of this kind, we performed a further categorisation exercise. Definitions of success were classified using the criteria for SMART goals (specific, measurable, ambitious, reasonable and time-bound) (Maxwell et al., 2015). We determined that whether goals were 'ambitious' and 'reasonable' was related to aspects of each organisation outside the scope of this review, therefore, each definition was assigned a combination of S, M and T depending on whether the information provided described a definition of success that was specific and / or measurable and / or time bound. If none of the criteria were met (or no definition was provided) 'NO' was recorded.

**Specific.** We looked for specificity in definitions of success, generally in terms of numbers (area of land, number of individuals, species etc). Specific definitions therefore tended to be measurable (but not always, for example, a definition that an organisation would achieve success by purchasing land to protect habitats was measurable but not specific). Terms such as 'improve', 'reduce', or 'increase' were not considered specific (but 'increase' would be measurable). Other terms not considered specific included 'enabling', 'maintaining', 'engaging', 'contributing', and 'facilitating' unless additional detail was provided.

**Measurable.** We looked for definitions of success that could be measured to determine whether or not they had been met. Specific goals tended to be measurable. Terms such as 'wilder' were not considered measurable, but 'increasing' or 'decreasing' were.

**Time-bound.** Definitions of success were assigned time-bound if they included a review date or clear time limit. We also included time considerations such as generational periods, for example, success defined as the second-generation of released species producing their own offspring. Mention of 'short-' or 'long-term' were not sufficient. Time bound was assumed if the objectives listed were contained within a time-bound larger document (e.g. strategy to 2027).

As with assigning the categories for definitions of success, if email / interview responses and online documents provided different details, the sum of all available data was used to assign SMT categories. Similarly, as we followed a generally conservative approach throughout, we erred on the side of assigning criteria, so if only one of several objectives was specific, measurable, and time-bound, the organisation was assigned SMT.

**Table S1.6 Most definitions of success used by rewilding organisations are related to social aims and objectives.** Definitions of success were extracted from rewilding organisations (by direct contact and from organisational documents) and coded according to 8 broad categories: social, knowledge sharing, ecosystem, species, diversity, habitat structure, abiotic, and other. Duplicates have been removed, and symbols such as '#' or words like 'species' used in place of specifics to make capture of repeats easier. Because by far the most mentions were of definitions of success related to the social category, a *double column* is provided to reduce the overall length of this table.

| Social | | Knowledge sharing | Ecosystem | Species | Diversity | Habitat structure | Abiotic | Other |
|---|---|---|---|---|---|---|---|---|
| Communities are empowered | Develop high-quality tourist infrastructure | Internationally recognized centre of excellence | Ecosystems are resilient | Released cheetahs survive independently | Diversity flourishes | Improve management of # area | Reduce emissions and increase sequestration of carbon | **Financial** – secure infrastructure, human and financial resources to achieve goals |
| Communities embrace conservation…emerging as stewards and champions | Public access to protected lands | Contribute to rewilding protocol | Resilient habitats | Improved knowledge of species' habitats and threats | Recruitment of native woody species | Plant # trees | | **Financial** – become sustainably profitable |
| Meaningful change to the benefit of businesses and people | Strengthen community connection with nature | Facilitate research | Sustain beauty and ecological integrity | Interventions to reduce threats to species and habitats | Recruitment of native non-woody species | Restore # area | | **Financial** – work on a range of financial mechanisms |
| Looking after heritage, people and natural resources | Engage and inspire communities | Establish botanical documentation and ecological research centre | Restore original ecology | Thriving species | Record new bird species | Secure # area of functional transboundary landscapes | | **Financial** – raise $ annually to support work |
| Enabling partnerships and improving livelihoods | Increase community agency | Develop effective and scientifically sound methodologies | Conserve at-risk ecosystems | # trees grown in nursery | Record new amphibian species | Create # natural assets | | **Financial** – ensure we build a sufficient capital base |

| | | | | | | | | |
|---|---|---|---|---|---|---|---|---|
| Empower local community members | Engage with local stakeholders and communities | Develop and demonstrate replicable and sustainable models that could be scaled up by others | Climate change mitigation | Contribute to genetic diversification | Areas of high diversity | Prevent degradation and encroachment onto hinterland | | **Targets** – contribute to national and international goals, priorities and targets |
| Find ways for humans and elephants to thrive together | Maintain acceptance of human communities toward species | Conservation education and awareness | Restore habitats for wildlife | Find ways for humans and elephants to thrive together | Propagate plant species | Bring areas under restoration efforts | | **Targets** – help deliver to "30% by 2030" global biodiversity framework goal |
| People and nature thrive | Training, research and employment opportunities | Produce reports, research papers and popular articles | Recover ecosystem functions | Future for elephants | Protect and preserve mammals, birds and amphibians | Land purchase | | **Legal** – Formal protection of priority habitats |
| Promote man's delight | Inspire and empower landowners | Support research projects for evidence based solutions | Protect ecological processes | Deliver programming for priority species | Biodiversity enhancement | Create rewilding areas across % land | | **Legal** – undertake legal actions necessary |
| Increase community engagement | Delivering significant benefits for people – healthier air, water and soils…health and wellbeing | Disseminate scientific data | Promote long-term survival of ecosystems | Reintroduce and propagate endangered flora | Protect biodiversity | Increasing amount of land | | **Legal** – defend, in court and out of court, the diffuse, collective and homogeneous individual rights and interests of the needy communities |
| Support # individuals within the community boosting income and well-being | Capturing hearts and minds…influence target groups | promote, support, disseminate, coordinate, develop and execute studies and research | Promote restoration of ecological interactions | Carry out translocation projects | Promote long-term survival of biodiversity | Restoring and reconnecting | | **Legal** – increasing legal protection |
| Increase household incomes | New sources of income for local communities | provide and perform services, advisory services and consultancy | Counter climate crisis | Counter species extinction crisis | Conserve significant biodiversity | Vast landscape…three untamed rivers | | **Legal** –assemble evidence required to submit a licence application |
| Build capabilities within communities | Build nature- and culture-based economy | Encourage research | Build environmental resilience to climate change | Increase viability of threatened species | Restore wildlife populations | Mosaic of habitats | | **Legal** – enable official designation |

| | | | | | | | | |
|---|---|---|---|---|---|---|---|---|
| Address socio-economic challenges | We support new businesses based on wild nature | Enhancing scientific education and training | Restore areas affected by human activities… complete and self-sustaining | Facilitate restoration of beavers | Increase diversity | Land protected from fragmentation / conversion / development / extractive uses | | **Legal** – lobby state, county and local government agencies |
| Solutions for harmonious coexistence of human beings and wildlife | Deliver positive ecological and socio-economic outcomes | Mobilise widespread public understanding | Restore extensive, connected, resilient landscape | First healthy wild-born juveniles | Abundant wildlife | Reconnect | | **Legal** – become official designated protected areas |
| Promote sustainable livelihood alternatives | Identify socio-economic barriers | Develop national strategies and policies | Restoration / re-establishment of natural processes | Increasing populations of keystone species | Overall uplift in biodiversity | Create corridors | | |
| Programme to ensure disease-free livestock | Grow network of young people | Develop a blueprint | Restore and reinstate as wide a range of natural processes as possible | Improve prey base for lynx, wolf etc | Inspiring, diverse mosaic of habitats and species | Provide habitat | | |
| Promote sustainable eco-development | Every young person no more than 50km from nature recovery project | Develop tools and platforms to share knowledge | Enhancing overall ecological health | Charismatic species | Discover and document biodiversity | Restore at least 50,000 hectares of critical ecosystems (forests, wetlands, savannas) | | |
| Generate sustainable income for local communities | Increase localism, resilience and self-sufficiency | Inspire and empower others to adopt and apply rewilding | Trophic chains | Replace non-native herbivores with those evolved with the native flora | Halt biodiversity decline | | | |
| Biodiversity friendly organic cocoa production | Break down social, cultural and political barriers | Demonstrate new models of land management…show case successes | Restore functionality | Establish a seed nursery to grow native plants and trees | | | | |
| Develop and implement community outreach program | Advocacy, education, and stewardship | Mainstream concept among decision makers | Support natural processes | Evaluating current conservation status | | | | |
| Educate and train people | Ensure cultures and systems support nature conservation | Provide sound scientific evidence for designing conservation-based initiatives | Support key habitats | | | | | |
| Generate opportunities and promote sustainable socioeconomic development | Strengthen and grow relationships with partners | Promote ecological science that is equitable and just | Provide sustainable water access for wildlife | | | | | |

| | | | |
|---|---|---|---|
| Projects and activities…education, culture, history, tradition, socio-economics, sports, tourism, conservation and preservation of the environment | Improve access to carbon funding for small-medium sized landholders | Be an educational resource | Conservation of watersheds and riparian ecosystems → resilience |
| Environmental education, ecotourism, socio-environmental public policies | Improve living conditions | Promote scientific research and indigenous knowledge on biodiversity, including the documentation of species, habitats, and local conservation practices | |
| Promotion of gender and peace | | Be a reference institution | |
| | | Collaborative alliances | |

#### 2.2.6 Monitor

'YES' or 'NO' was recorded in this column to understand the number of organisations providing detail on how they monitor progress.

#### 2.2.7 Monitor direct extract and classification

Details of how organisations monitor progress in rewilding were taken directly from documents, websites, or direct communication with GRA organisations. Within documents, some organisations clearly outlined their monitoring activities. If clear monitoring strategies were not provided, we searched within documents for "monitor", "measure", and "track". Similar to the grey literature on definitions of success, this data was often long-form, detailed and not easy to compare across organisations. Extracted information was coded into seven broad categories (Table S1.7): fieldwork, remote sensing, wildlife tracking, DNA methods, consultation, experiment, and aspirational (where documents or contacts mentioned plans for future monitoring). More than one category was assigned where appropriate. If there were different responses in contact and documents, all were included for categorisation.

These categories are close to those used for investigating monitoring methods in the academic literature, and definitions of each category are the same as above. The review category was removed for grey literature as none of the organisations made use of review to monitor progress, and an aspirational category was added to capture organisations' plans to introduce or implement monitoring methods in the future. Aspiration was assigned alongside other categories if there were plans for monitoring in addition to current activities.

**Table S1.7 Fieldwork is the most common broad method used to monitor progress in rewilding organisations.** Methods used to monitor progress were extracted from rewilding organisations (by direct contact and from organisational documents) and coded according to 7 broad categories: fieldwork, remote sensing, wildlife tracking, DNA methods, consultation, experiment, and aspirational. Duplicates have been removed, and words like 'species' used in place of specifics to make capture of repeats easier.

| Fieldwork | Wildlife tracking | Remote sensing | DNA methods | Consultation | Experiment | Aspirational |
|---|---|---|---|---|---|---|
| Field observations | GPS collar | Tree mapping | Genetic tracking | Solicit feedback | Exclosure plots | Genetic monitoring |
| Ecological baseline surveys | Camera trapping | Satellite imagery | Detect DNA in water samples | Social studies | Different herbivore mixes – compare with baseline | eDNA |
| Species counts | Acoustic monitoring | GPS mapping | DNA barcoding | Community workshops | | 3D modelling |
| Driving surveys | Autonomous camera drones | Lidar | DNA metabarcoding | Monitoring Groups were set up for collaboration and joint cocreation | | AI camera tracking |
| Foot patrols | Radio telemetry | Drones to produce maps | Soil eDNA | | | Satellite imagery |
| Waterhole surveys | Transmitters | NDVI vegetation condition mapping | Scientific breeding for conservation | | | Remote sensing |
| Hilltop surveys | Direct observation | | | | | Field surveys |
| Vegetation transects | | | | | | GIS |
| Livestock counts | | | | | | Bioacoustics |
| Tree measurements | | | | | | AI-based biodiversity |

| | | | | | | |
|---|---|---|---|---|---|---|
| | | | | | | monitoring cameras |
| Daily infrastructure monitoring | | | | | | Remote monitoring equipment |
| Aerial census | | | | | | Long-term soil sampling – soil chemistry |
| Biodiversity assessments | | | | | | Additional fieldwork |
| Tree enumeration | | | | | | Establish monitoring plan / strategy |
| Plant species ID | | | | | | Additional wildlife tracking |
| Routine monitoring | | | | | | Consultation |
| River surveys | | | | | | |
| Forest surveys | | | | | | |
| Monthly census | | | | | | |
| Mark and release | | | | | | |
| Habitat classification | | | | | | |
| Malaise traps | | | | | | |
| Transects | | | | | | |
| Browsing impact assessments | | | | | | |
| Long-term monitoring plots | | | | | | |
| Fixed point photographs | | | | | | |
| Scientific expeditions | | | | | | |

#### 2.2.8 Evaluate

'YES' or 'NO' was recorded in this column to understand the number of organisations providing detail on how they evaluate success.

#### 2.2.9 Evaluate direct extract and classification

We wanted to understand the kinds of metrics used by organisations to evaluate success in rewilding. We took information taken directly from documents, websites, or direct communication with GRA organisations. Within documents, extraction was typically from impact statements or project summaries. If clear evaluation measures were not provided, we searched within documents for "evaluate" and "assess". Again, this data was often long-form and difficult to compare so it was classified into nine broad categories: internal standards, social, species, academic, diversity, financial, area, external standards, and aspirational. More than one category was assigned where appropriate. If there were different responses in contact and documents, all were included for categorisation.

> **Internal standards.** This was a broad category that captured metrics or measures that were mostly relevant within the GRA organisation itself. It includes number of internal personnel hired or trained, size of organisation, number of projects planned and implemented etc. If counts were used as indicators of success outside of any context linking them to diversity of species – number of trees planted, for example – then this was counted as an internal standard. If an organisation listed, for example, number of *native* trees planted or grown, then this would be categorised under diversity. We also grouped measures of engagement, number of visitors etc in this category, as they are measures by which organisations measure their success through rather than the impact of those engagements. This category also captured organisations that have developed their own internal evaluation strategies.
>
> **Social.** Many organisations used social measures to evaluate their progress and success in rewilding. These measures include training and education programmes designed and delivered, funds distributed to communities, jobs created, wellbeing assessment and collaborative activities. It also captured

measures related to non-wild animals (e.g. livestock health and management programmes).

**Species.** Organisations focused on threatened, reintroduced, or charismatic species typically used species-centric measures to evaluate success, such as number of individuals, survival rate, reproductive outcomes etc.

**Academic**. Some organisations measured success by the extent to which they were able to disseminate information. This category includes number of academic publications, other articles, data sets etc shared, as well presentations at academic events and promotion of their works within the academic and practitioner communities.

**Diversity.** This group was for organisations evaluating success through records of numerous or new mammal, bird, plant, fungi etc species.

**Financial.** This category captured organisations evaluating success linked to their own financial success, for example, measures of money raised, money allocated, income, grants secured etc.

**Area.** Several organisations evaluated success by the size or number of areas bought, protected, conserved, restored etc.

**External standards.** Some organisations used standards set by external bodies or organisations to evaluate their progress. To be counted in this category, it had to be clear that the GRA organisation measured their own success with reference to such external standards. For example, several organisations described themselves as 'aligned' with or to several of the sustainable development goals. These were only assigned to the external standards category, though, if additional context showed that meeting that alignment was an evaluation concern of the organisation. For example, some organisations include sustainable development goals alignment as part of their own broader metrics for evaluation.

**Aspirational.** This category was used to capture planned or envisioned evaluation activities that organisations described.

**Table S1.8 Measures and standards for evaluation used by rewilding organisations coded to 9 broad categories: internal standards, social, species, academic, diversity, financial, area, external standards, and aspirational.** Measures for evaluating success were extracted from rewilding organisations (by direct contact and from organisational documents). Duplicates have been removed, and symbols such as '#' or words like 'species' used in place of specifics to make capture of repeats easier.

| Internal standards | Social | Species | Academic | Diversity | Financial | Area | External standards | Aspirational |
|---|---|---|---|---|---|---|---|---|
| Numbers of employees / volunteers | Jobs created | Reproductive outcomes | Research outputs | Mammal species found | Number of donors | Number of protected areas | Eight Pillars of Positive Peace | Volunteer events |
| Number of countries worked in | People living in landscapes | Target populations | Publications | Bird species found | Increase in investment | Size of protected area(s) | Sustainable Development Goals alignment | News articles |
| Number of organisations / partnerships | Human wellbeing | Tree survival rate | Technical reports | Wildlife populations | Income | Connectivity restored | EU Biodiversity strategy | Reintroduction |
| Households surveyed | Livestock care | Number of target / focal species | Data sets shared | Species counts | Carbon credits | Forest cover area | IUCN Marseille Resolution 127 | Trial |
| Baseline surveys done | Training | Unique species | Citations | # native species grown | Financial metrics | Creation of parks / protected areas | Aichi target 11 | Long term case studies |
| Projects planned | Education | Tracking collars deployed | Scientific cooperation project | # threatened species grown | Grants secured | Growth of protected areas | IUCN species of concern | Board review progress |
| Projects implemented | Money distributed | Reintroductions | Citizen science observations | Species diversity | Use of contract £ | | | Undefined |

| | | | | | | | | |
|---|---|---|---|---|---|---|---|---|
| Natural assets | Scholarships | Recovery of focal species | Presentations | Vertebrate species recorded | £ raised | | | Increase area |
| Governance index | Community revenue | Species released | COP15 | Fungi species recorded | £ allocated | | | Photo monitoring for evaluation |
| Programmes run | People impacted | Species tagged | Promotion of definitions and methods | Unique plant and animal species | | | | |
| Equipment | Economic empowerment | Species abundance | | | | | | |
| # visitors | Health | GPS marked birds | | | | | | |
| # engagements | Gender inclusion | Translocations | | | | | | |
| Amount of planting done | Social innovation | Recolonisations | | | | | | |
| Donations | Social collaboration | Reports of focal species | | | | | | |
| Surveys | Donations | Species benefits | | | | | | |
| Set of evaluation questions | Jobs | Species colony health | | | | | | |
| Online followers | Social metrics | | | | | | | |
| Recognition of internal criteria | Loans | | | | | | | |
| Lobbying | Business networks | | | | | | | |
| Setting internal criteria | £ spent with local suppliers | | | | | | | |

| | |
|---|---|
| # members | Supporting indigenous-led conservation |
| Convening stakeholders | Feral animal control |
