## Supplementary 2 - extended results for "Systematic mapping shows monitoring, evaluation, and engagement are needed to build an evidence base for rewilding"

This supplementary information contains extended results for the systematic map:

### Extended Results

This section of supplementary information contains additional context:

- Lists of academic and grey literature components of this systematic map
- Supporting results figures

### 1. Results of academic and grey literature searches

Our academic literature review comprised a full review of 48 papers (Table S2.1). We included 93 partner organisations from the Global Rewilding Alliance (GRA)

**Table S2.1 List of academic papers included for academic literature component of systematic map.**

| | year | title | reference |
|---|---|---|---|
| **1** | 2022 | Monitoring rewilding from space: The Knepp estate as a case study | Schulte to Bühne et al. (2022) |
| **2** | 2022 | Expert-based assessment of rewilding indicates progress at site-level, yet challenges for upscaling | Segar et al. (2022) |
| **3** | 2021 | Rewilding and gazetting the Ibera National Park: Using an asset approach to evaluate project success | Pettersson and de Carvalho (2021) |
| **4** | 2018 | Measuring rewilding progress | Torres et al. (2018) |
| **5** | 2022 | Long-term effects of rewilding on species composition: 22 years of raptor monitoring in the Chernobyl Exclusion Zone | Dombrovski et al. (2022) |
| **6** | 2021 | Enhancing monitoring of rewilding progress through wildlife tracking and remote sensing | Mata et al. (2021) |
| **7** | 2023 | Rewilding giant tortoises engineers plant communities at local to landscape scales | Tapia and Gibbs (2023) |
| **8** | 2023 | Long-term monitoring of mammal communities in the Peneda-Gerês National Park using camera-trap data | Zuleger et al. (2023) |
| **9** | 2019 | Using lidar to assess the development of structural diversity in forests undergoing passive rewilding in temperate Northern Europe | Thers et al. (2019) |
| **10** | 2018 | From exploration to establishment: Activity changes of the first collared peccary (Pecan tajacu) group reintroduced in South America | Hurtado et al. (2018) |
| **11** | 2024 | Long-term vegetation responses to climate depend on the distinctive roles of rewilding and traditional grazing systems | Rincon-Madroñero et al. (2024) |
| **12** | 2019 | Implications of Spatial Habitat Diversity on Diet Selection of European Bison and Przewalski's Horses in a Rewilding Area | Zielke et al. (2019) |
| **13** | 2018 | Rewilding cultural landscape potentially puts both avian diversity and endemism at risk: A Tibetan Plateau case study | Li et al. (2018) |

| | | | |
|---|---|---|---|
| **14** | 2017 | Rewilding the Atlantic Forest: Restoring the fauna and ecological interactions of a protected area | Fernandez et al. (2017) |
| **15** | 2016 | Rewilding with large herbivores: Direct effects and edge effects of grazing refuges on plant and invertebrate communities | Van Klink et al. (2016) |
| **16** | 2016 | Rural abandoned landscapes and bird assemblages: winners and losers in the rewilding of a marginal mountain area (NW Spain) | Regos et al. (2016) |
| **17** | 2023 | Evaluating the performance of conservation translocations in large carnivores across the world | Thomas et al. (2023) |
| **18** | 2021 | Restoring a butterfly hot spot by large ungulates refaunation: the case of the Milovice military training range, Czech Republic | Konvička et al. (2021) |
| **19** | 2023 | Reintroducing bison to Banff National Park - an ecocultural case study | Heuer et al. (2023) |
| **20** | 2023 | Intensive agriculture as the main limiting factor of the otter’s return in southwest France | Couturier et al. (2023) |
| **21** | 2023 | Mini Safe Havens for population recovery and reintroductions 'beyond-the-fence' | Smith et al. (2023) |
| **22** | 2021 | Ten years on: have large carnivore reintroductions to the Eastern Cape Province, South Africa, worked? | Banasiak et al. (2021) |
| **23** | 2020 | Spatial patterns of the first groups of collared peccaries (Pecari tajacu) reintroduced in South America | Hurtado et al. (2020) |
| **24** | 2017 | Using ecosystem engineers as tools in habitat restoration and rewilding: beaver and wetlands | Law et al. (2017) |
| **25** | 2022 | Introduction of giant tortoises as a replacement ecosystem engineer to facilitate restoration of Santa Fe Island, Galapagos | Tapia et al. (2022) |
| **26** | 2022 | Active versus passive restoration: Forests in the southern Carpathian Mountains as a case study | Hartup et al. (2022) |
| **27** | 2017 | Refaunation and the reinstatement of the seed-dispersal function in Gorongosa National Park | Correia et al. (2017) |
| **28** | 2023 | Recolonizing native wildlife facilitates exotic plant invasion into Singapore's rain forests | Ho et al. (2023) |
| **29** | 2022 | Herbivore exclusion and active planting stimulate reed marsh development on a newly constructed archipelago | Temmink et al. (2022) |
| **30** | 2023 | Narratives of land abandonment in a biocultural landscape of Spain | Quintas-Soriano et al. (2023) |
| **31** | 2024 | Rewilding by large ungulates contributes to organic carbon storage in soils | Kaštovská et al. (2024) |

| | | | |
|---|---|---|---|
| **32** | 2024 | Long-term vegetation trajectories to inform nature recovery strategies: The Greater Côa Valley as a case study | Elphick et al. (2024) |
| **33** | 2015 | Rewilding: Pitfalls and opportunities for moths and butterflies | Merckx (2015) |
| **34** | 2024 | Evaluating the success of vegetation restoration in rewilded salt marshes | Carneiro et al. (2024) |
| **35** | 2020 | Assessing the effects of habitat management practices on vocalizing animals using passive acoustic monitoring.<br>Specifically: Effects of Different Habitats Created by Rewilding on Bat and Bird Activity and Diversity | Beason (2020) |
| **36** | 2021 | The response of plants, carabid beetles and birds to 30 years of native reforestation in the Scottish Highlands | Warner et al. (2021) |
| **37** | 2022 | Does restoring native forest restore ecosystem functioning? Evidence from a large-scale reforestation project in the Scottish Highlands | Warner et al. (2022) |
| **38** | 2024 | Monitoring the rewilding of the Brazilian Atlantic Forest on tree and mammal diversity: From a biodiversity hotspot to a biodiversity hopespot | Merelli et al. (2024) |
| **39** | 2024 | Time to independence and predator-prey relationships of wild-born, captive-raised cheetahs released into private reserves in Namibia | Marker et al. (2024) |
| **40** | 2024 | Rewilding landscapes with apex predators: cheetah (Acinonyx jubatus) movements reveal the importance of environmental and individual contexts | Dimbleby et al. (2024) |
| **41** | 2024 | Restoring ecological function: Interactions between vertebrates and latrines in a reintroduced population of Rhinoceros unicornis | Awasthi et al. (2024) |
| **42** | 2024 | Born to be wild: Captive-born and wild Iberian lynx (Lynx pardinus) reveal space-use similarities when reintroduced for species conservation concerns | Cisneros-Araujo et al. (2024) |
| **43** | 2024 | Population status and genetic assessment of mugger (Crocodylus palustris) in a tropical regulated river system in North India | Sharma et al. (2024) |
| **44** | 2022 | The impacts of reintroducing cheetahs on the vigilance behaviour of two naive prey species | Welch et al. (2022) |
| **45** | 2024 | Topographic heterogeneity influences diversity and abundance of Orthoptera in a rewilding scheme | Gardiner and Casey (2024) |
| **46** | 2019 | Bare soil cover and arbuscular mycorrhizal community in the first montane forest restoration in Central Argentina. | Becerra et al. (2019) |
| **47** | 2021 | Avifaunal responses after two decades of Polylepis forest restoration in central Argentina. | Barri et al. (2021) |
| **48** | 2022 | Recommendations for the rehabilitation and release of wild-born, captive-raised cheetahs: the importance of | Walker et al. (2022) |

| | | |
|---|---|---|
| | | pre-and post-release management for optimizing survival. |

**Table S2.2 List of Global Rewilding Alliance members included for grey literature component of systematic map**

| | Organisation |
|---|---|
| 1 | African Conservation Foundation |
| 2 | Cheetah Conservation Fund |
| 3 | Conserve Global |
| 4 | Endangered Wildlife Trust |
| 5 | Enonkishu Conservancy |
| 6 | Ferncliffe |
| 7 | Global Humane Conservation Fund of Africa |
| 8 | Mali Elephant Project |
| 9 | MKAAJI MPYA asbl |
| 10 | Peace Parks Foundation |
| 11 | Save The Elephants |
| 12 | Pakistan Environment Trust |
| 13 | Balipara Foundation |
| 14 | Chinese Felid Conservation Alliance |
| 15 | Edhkwehlynawd Botanical Refuge Centre Trust |
| 16 | Junglescapes |
| 17 | Junglo |
| 18 | Planet Indonesia |
| 19 | The Corbett Foundation |
| 20 | Wildlife Trust of India |
| 21 | AMAP Brazil |
| 22 | Araguia - Instituto Araguaia de Proecao Ambiental |
| 23 | Black Jaguar Foundation |
| 24 | Ecosistemas Argentinos |
| 25 | Felinos do Aguaí Institute |
| 26 | Guanacaste Dry Forest Conservation Fund |
| 27 | Instituto Fauna Brasil |

| | |
|---|---|
| **28** | Instituto Homem Pantaneiro |
| **29** | Instituto Serra do Tangará |
| **30** | Ofrenda A'bunna |
| **31** | OSA Conservation |
| **32** | Reforestamos |
| **33** | Rewilding Argentina |
| **34** | Rewilding Chile |
| **35** | Sociedale de Pesquisa em Vida Selvagem (SPVS) |
| **36** | Terra Habitus |
| **37** | Wildtracks |
| **38** | ARK Nature Foundation |
| **39** | Beaver Trust |
| **40** | Bunloit rewilding |
| **41** | Carpathia European Wilderness Reserve |
| **42** | Kent Wildlife Trust |
| **43** | Knepp Wildland Foundation |
| **44** | Nattergal |
| **45** | Rewilding Britain |
| **46** | Rewilding Europe |
| **47** | Rewilding Oder Delta |
| **48** | Rewilding Portugal |
| **49** | Rewilding Sweden |
| **50** | Scotland: the Big Picture |
| **51** | The European Nature Trust |
| **52** | Whole Wild World |
| **53** | Wild Europe |
| **54** | Wildwood Trust |
| **55** | Youngwilders |
| **56** | Conservation Beyond Borders |
| **57** | Mossy Earth |
| **58** | Re:wild |
| **59** | American Prairie Reserve |
| **60** | Canadian Parks and Wilderness Society, CPAWS |
| **61** | ICCF (International Conservation Caucus) Group |

| | |
|---|---|
| **62** | NorthEast Wilderness Trust |
| **63** | Rewild New Jersey Community Cooperative |
| **64** | Rewilding America Now |
| **65** | Southern Plains |
| **66** | Wyoming Wilderness Association |
| **67** | Yellowstone to Yukon Conservation Initiative |
| **68** | Bush Heritage Australia |
| **69** | Gondwana Link |
| **70** | Great Eastern Ranges |
| **71** | South Endeavour Trust |
| **72** | The Forktree Project |
| **73** | For The Love of Wildlife |
| **74** | Action for Protection of Wild Animals (APOWA) |
| **75** | Altyn Dala Conservation Initiative and ACBK - Association for the Conservation of Biodiversity of Kazakhstan |
| **76** | Raintree Foundation |
| **77** | Wildlife Conservation Trust |
| **78** | Citizen Zoo - Rewilding our Future |
| **79** | Climate Action North |
| **80** | Deutsche Umwelthilfe |
| **81** | Embercombe retreat centre |
| **82** | Irish Trees |
| **83** | Trees For Life |
| **84** | Kentucky Natural Lands Trust |
| **85** | Re-wilding Initiative |

### 2. Supporting figures for results

#### 2.1 Case study locations.

Our systematic map found case studies in the academic and grey literature around the world (Figure S2.1).

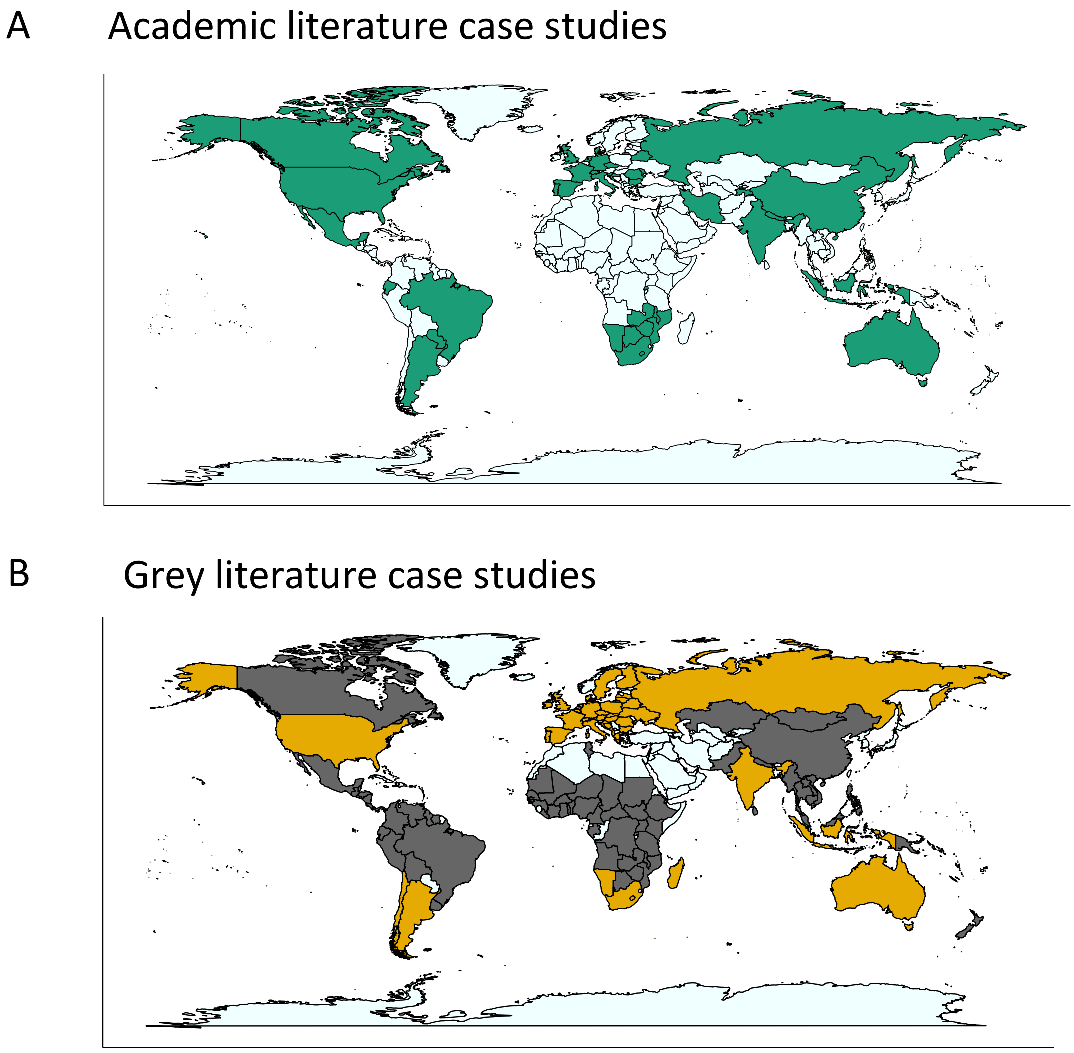

**Figure S2.1 Rewilding is happening globally.** Academic rewilding case studies were described in 40 countries, and the members of the Global Rewilding Alliance (GRA) included in this review have rewilding case studies in 106 countries. **A.** Countries in which academic rewilding case studies were described are coloured green. The median number of case studies in each country was 1 (interquartile range (IQR) = 1). In some instances, a single paper reviewed multiple case studies in a single country. For example, the most case studies (n = 9) were described in Czech Republic, but 8 of these were in a single paper, Kaštovská et al. (2024).

Thomas et al. (2023) described 33 case studies across 21 countries. Inset shows the number of studies describing monitoring only, evaluation only, or both monitoring and evaluation. **B.** The countries in which GRA members have rewilding case studies are coloured: all shaded countries – grey and yellow – represent the total 106 countries. Countries shaded yellow are those where an organisation responded to direct communication by email. The median frequency for each country was 1 (IQR = 1), with most organisations operating in the UK (n = 18), the USA (n = 12), and India (n = 10). Countries were extracted directly from the GRA partner organisation details provided online. Inset shows the number of organisations that provided information (or for which information could be found) on their definitions of success, use of monitoring, and evaluation.

#### 2.2 Case study area

More very large rewilding areas were described in the grey literature than in the academic. 29 academic papers (60%) were explicit about the size of rewilding area included in case studies and most covered relatively small study areas (Figure S2.2). The median area of academic case study described was 13.7km$^2$ (interquartile range (IQR) = 57.2km$^2$). In the grey literature, we were able to determine the size of rewilding project for 56 organisations using email responses and the grey literature available (66% of all projects included). The median area of rewilding case study described in the grey literature was 309km$^2$ (IQR = 9756km$^2$).

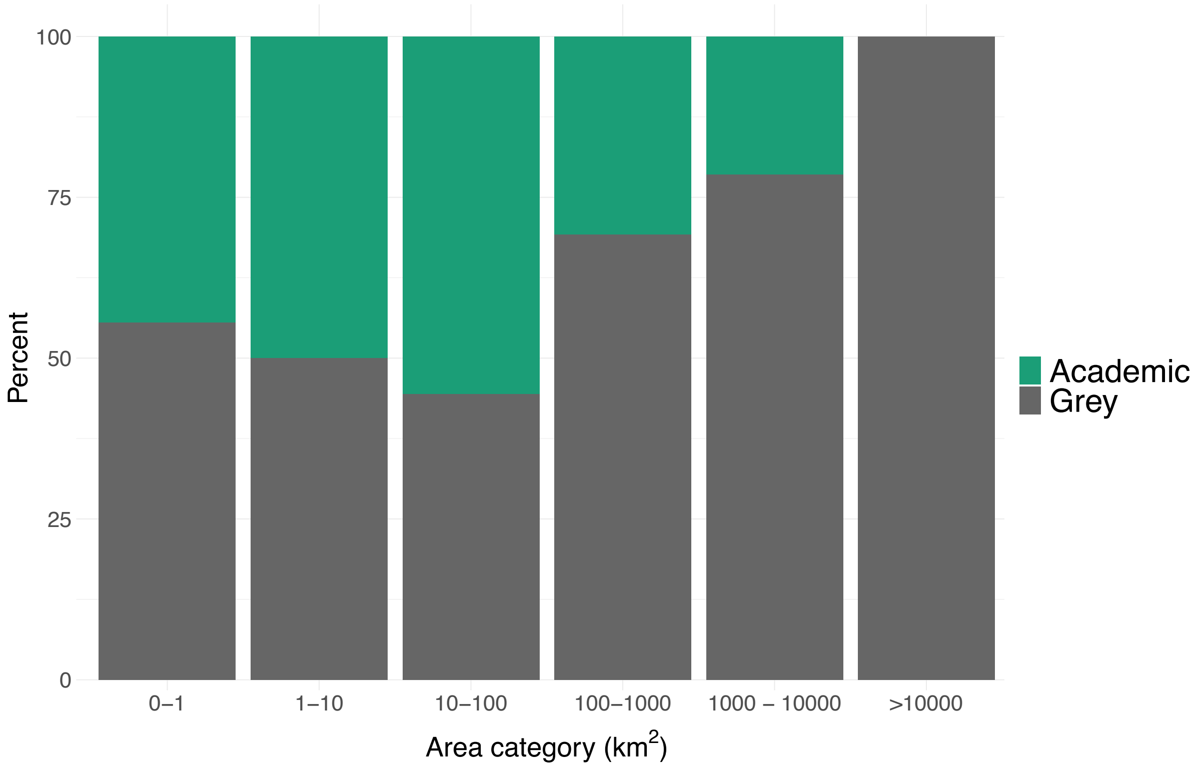

**Figure S2.2 More very large rewilding areas were described in the grey literature than in the academic.** Academic case studies and GRA projects were grouped according to the area of land they included across 6 categories: up to 1km$^2$, 1-10 km$^2$, 10-100 km$^2$, 100-1000km$^2$, 1000-10,000 km$^2$, and >10,000 km$^2$. Academic articles were only included where they were clear about the area of land involved in the study (n = 29 out of 48 studies). It is important to note that direct comparison is not simple here, as many academic case studies described small patches, experimental areas, or individual habitat areas within much larger rewilding sites.

### 2.3 Case study duration

Case studies in the academic literature typically cover short time periods (Figure S2.3A). We compare this with the duration of operation for rewilding organisations (Figure S2.3B). We acknowledge that these data reflect different things – the length of time a rewilding organisation has been operational is not necessarily linked to the length of time rewilding has been implemented across their whole area, and academic studies will almost always have to report on a shorter period within a rewilding project's progress – however, this disparity is worthy of record. Rewilding organisations will have long-term perspectives and will likely want to take evidence-based policy decisions. But if most of the evidence currently being offered is based on narrow study snapshots, it may not offer the best possible guidance.

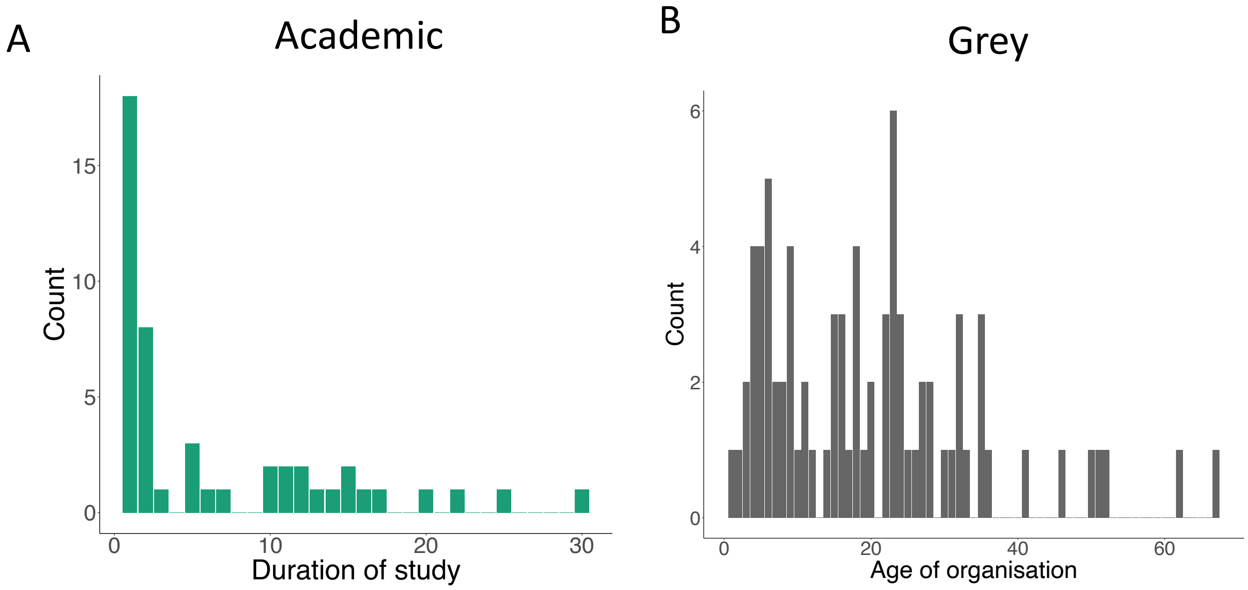

**Figure S2.3 Academic case studies tend to describe short time periods and small numbers of sites. A.** Most academic case studies reported on one or two years of data (26 out of 48 studies, median duration = 1 year, IQR = 1). The longest duration of study was 30 years (Rincon-Madroñero et al., 2024). **B.** As an indirect comparison, some rewilding organisations have been established for several decades. Where organisations provided a date of foundation / establishment, this was subtracted from 2025 to give the age of organisation. We were able to include the duration of operation for 79 rewilding organisations. The longest-running organisation had been running for 67 years and the shortest just one year.
